# Identification of disease-specific vulnerability states at the single-cell level

**DOI:** 10.1101/2024.12.04.626873

**Authors:** Matthew D’Antuono, Madison Sharp, Rishika Chowdary, Michael E. Ivan, Ricardo J. Komotar, Robert K. Suter, Nagi G. Ayad

**Author notes:** Contact Information: Nagi G. Ayad, Ph.D. (Lead Contact) Georgetown University, Department of Oncology 3970 Reservoir Rd NW, Washington, D.C. 20057,; Robert K. Suter, Ph.D., Georgetown University, Department of Oncology 3970 Reservoir Rd NW, Washington, D.C. 20057.

## Abstract

Intratumor heterogeneity in glioblastoma (GBM) impedes successful treatment as it is not obvious which tumor cells should be targeted. Here, we posit that single-cell-resolution transcriptomic data can be integrated with loss-of-function screens to identify the most critical cells to target within a tumor. We parsed CRISPR screen data from the Dependency Map (DepMap) Consortium and identified a GBM Dependency Signature (GDS) − 168 genes that are essential for GBM cell viability in vitro. Through similarity scoring of GDS transcriptomic profiles in single-cell RNA-sequencing (scRNA-seq) data and iterative hierarchical clustering, we identify and report 3 single-cell vulnerability states (VS) characterized in 49 GBM tumors using both scRNA-seq and spatial transcriptomic data. These VS reflect single-cell gene dependencies and differ significantly in enrichment profiles and spatial distributions. Additionally, each VS is differently sensitive to cancer drugs, with VS2 solely responsive to temozolomide treatment. Importantly, the proportion of VS in each GBM tumor is variable, suggesting a means of stratifying patients in clinical trials. Collectively, we have developed a novel computational pipeline to identify unique vulnerability states in GBM and other cancers, which can be used to identify existing or novel drugs for incurable diseases.

## INTRODUCTION

Glioblastoma (GBM) is the most common primary malignant brain tumor in adults. Development of effective treatments for GBM is challenging, and despite extensive research efforts no new therapies have been approved for GBM since the establishment of the current standard-of-care nearly two decades ago.^1^ Without advancements in treatment for GBM, the prognosis is poor with a median overall survival after diagnosis of only 15 months and a 5-year survival rate less than 5%.^1–3^ Therefore, new GBM treatments are desperately needed.

One of the driving forces behind treatment and drug development challenges is the extensive inter- and intra-tumor heterogeneity innate to GBM. Intratumor heterogeneity (ITH), in particular, presents a large hurdle. For example, different subclonal sensitivities to EGFR inhibitors in GBM have been observed *in vitro* and *in vivo* with additional subclone-level differences noted in GBM progression, invasion, and animal survival.^4^ Each patient tumor contains cells with high degrees of genomic, transcriptomic, proteomic, and metabolic variability, and this can lead to innate or rapid development of drug resistance and subsequent treatment failure.^5–10^

To date, much of the heterogeneity observed in GBM is the result of recurrent transcriptional modules expressed during cellular development and differentiation or in response to stressful stimuli such as hypoxia.^8,11–16^ In GBM, tumor cells exist along a spectrum of transcriptional cell states that mimic canonical neurodevelopmental cell types.^8^ These states are designated as astrocyte-like (AC), mesenchymal-like (MES), neural progenitor-like (NPC), and oligodendrocyte-like (OPC).^8^ Multiple studies have demonstrated that these states are highly plastic and the relative proportions of each in a tumor will change under treatment pressure, contributing to drug resistance and tumor recurrence.^8,13,14,17–19^

As GBM heterogeneity continues to impede drug discovery, a novel method is needed to identify the distinct transcriptional cell states to target to achieve therapeutic efficacy. This framework involves prioritization of those states identified as most essential to a tumor with simultaneous targeting of the remaining cancer cells. Toward that end, we hypothesized that single-cell transcriptomic data could be integrated with loss-of-function screens to identify the most essential cells within a GBM tumor.

Genome-wide loss-of-function screens such as those conducted by the Dependency Map (DepMap) Consortium offer insight into genetic vulnerabilities of cancer cell lines and make for promising resources to identify candidate genes for targeted therapy development. DepMap performed systematic CRISPR knockout screens in more than 1,000 cell lines across nearly every type of cancer lineage, creating an unparalleled collection of genetic profiles that quantify the essentiality of nearly every gene to each cell lines’ viability.^20–24^ Importantly, DepMap data has already been successfully leveraged to discover genetic vulnerabilities in cancer cells with translation to *in vivo* models and patient tumors.^25–28^

We sought to leverage CRISPR gene effect scores (ES) from DepMap to identify exploitable “vulnerability states” in GBM – single-cell states with shared, therapeutically-relevant transcriptional profiles that would be useful toward informing drug development strategies. We identified a list of GBM lineage-specific essential genes and used this gene set to inform transcriptomic single-cell state assignments via similarity scoring and iterative hierarchical clustering. We uncovered three (3) novel vulnerability states within and across GBM tumors from different datasets that express GBM dependency genes to varying degrees. We found these states to have distinct enrichment profiles in both single-cell RNA-seq and spatial transcriptomic data, and they are each spatially distinct among patient GBM tumor sections. Critically, these states are distinct from existing GBM transcriptional classifiers and are informed by CRISPR screen-derived, lineage-specific essential genes, paving the way for drug discovery and development efforts using these single-cell states. Moreover, each VS has distinct functional enrichments and each is differently sensitive to FDA-approved cancer therapies. Specifically, we show that one cell state is uniquely sensitive to temozolomide (TMZ) chemotherapy while the other two are resistant, and this resistance is only overcome by combination therapy. The presented methodology and framework for identification of dependency signature-derived cell states, therefore, has far reaching implications for personalized medicine in GBM and could be applied to other types of cancer.

## RESULTS

### Analysis of DepMap genome-wide knockout screen uncovers 168-gene glioblastoma dependency signature (GDS)

We acquired DepMap’s Project Achilles open-access genome-wide CRISPR knockout screen post-Chronos effect score (ES) data from their online portal. Lower ES for a gene indicates a cell line is more dependent on that gene for survival. In general, an ES of -1 is commensurate with the median of all common essential genes, while an ES of 0 implies that a gene is not essential to a cell line. The DepMap ES dataset (23Q2 release) contains ES for 17,453 genes across 1,078 cell lines (Figure 1A). We interrogated these data to identify genes whose knockdown negatively impacts viability in GBM cell lines specifically. In our approach, we strived to select for genes uniquely essential to GBM over other cell line lineages with the aim of identifying genes that, if targeted for knockdown/out, would have minimal off-target/tumor effects and thereby minimize potential toxicities. Moreover, through these methods we sought to minimize identification of pan-cancer essential genes (e.g. CDK1, DNMT1, HDAC3) whose inhibitors have suffered from clinical trial failures due to on-and off-target toxicities.

**Figure 1:**
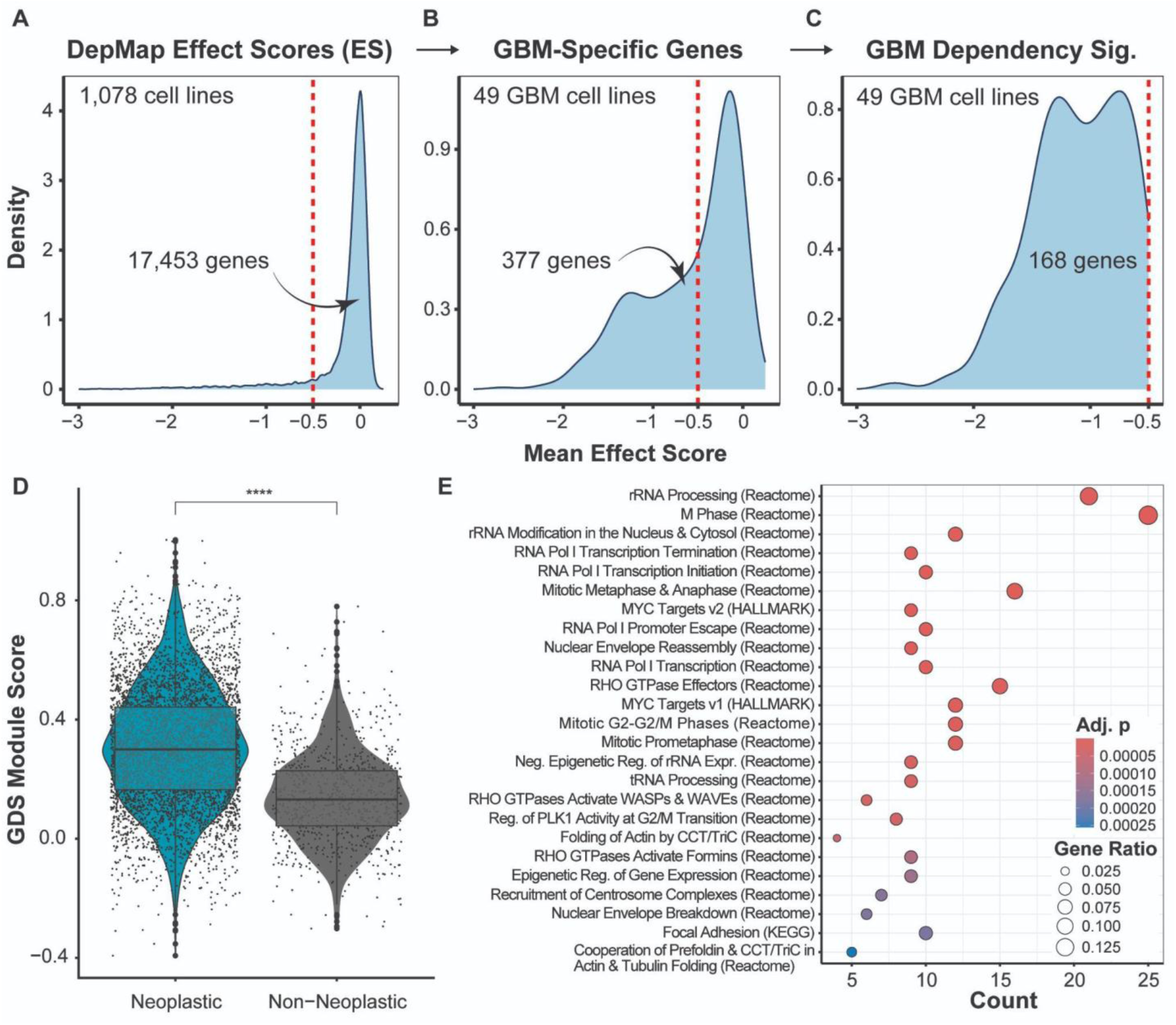
Analysis of DepMap genome-wide knockout screen uncovers 168-gene glioblastoma dependency signature. (**A-C**) Density plots depicting the selection pipeline for the Glioblastoma (GBM) Dependency Signature (GDS). The red dashed line in each represents a mean effect score of -0.5. (**A**) shows the mean effect scores (ES) of all genes (n = 17,453) in all cell lines (n = 1,078) present in the DepMap dataset. After applying a linear model comparing ES in GBM-lineage cell lines (n = 49) to all other cell lines, (**B**) depicts the mean ES of 377 GBM-specific genes. (**C**) then represents the mean ES of the 168-gene GDS, identified by filtering the 377 genes from (**B**) for those with mean ES < -0.5. (**D**) Violin and overlapping box plot depicting the module scores (MS) of the GDS in each single-cell (black dots) in the Neftel et al. scRNA-seq dataset (5,689 cells, 20 adult GBM tumors), grouped by neoplastic status. Neoplastic cells (n = 4,865) are compared to non-neoplastic cells (n = 824, consisting of T-cells, macrophages, and oligodendrocytes) by a Student’s t test (**** = p < 0.0001). (**E**) Dot plot representing Molecular Signatures Database (MSigDB) over-representation analysis performed on the 168-gene GDS (using collections C2 and H). The top 25 over-represented terms here have Benjamini-Hochberg (BH)-adjusted p-value < 0.05 by Fisher’s exact test. Terms (pathways) are sorted by descending adjusted p-value (bar color). The gene ratio is the number of GDS genes that are enriched in a pathway over the total number of genes that define it.

With these goals in mind, we compared ES for every gene in GBM cell lines to the corresponding ES in all other cell line lineages using a generalized linear model (GLM). Of 1,078 cell lines in the dataset, 49 are GBM-lineage specific (Table S1). Genes with significantly lower ES in GBM lines were defined as those with negative effect sizes and a p-value < 0.05 by GLM (Figure 1B). Using these criteria, we identified 377 genes whose knockout more significantly impacted GBM cell line viability than other cell lineages (Table S2). Importantly, a strict ES cutoff was then set to identify which of the 377 genes kill GBM cells *in vitro* – removing genes with mean ES > 0, for example. We imposed a cutoff of mean ES < -0.5 in GBM cell lines as this quantitative value has been used in previous studies and purported as representing depletion in most cell lines.^23,27^ We thereby identified 168 genes uniquely essential to GBM cell lines, a list we term a GBM Dependency Signature (GDS) (Figure 1C, Table S3). Some of the top GDS genes by GLM-derived p-value include PRKAR1A, LMNA, JUN, VRK1, MTOR, RAC1, CDC42, and CDK2.

To confirm the specificity of our analysis and of the resulting essential genes to GDS, we performed the same analysis on BC, PDAC, and multiple myeloma (MM) cell lines in the DepMap dataset. There were 132 BC-specific essential genes, 115 PDAC-specific essential genes, and 274 MM-specific essential genes. Importantly, the majority of each of these gene sets were unique to each cancer lineage when compared to the GDS, with minimal overlap across cancer lineages. Interestingly, there were 0 overlapping genes between any three of these gene sets, emphasizing the specificity of our approach (Figure S1D).

To determine GDS gene expression in patient GBM tumors, we used a single-cell RNA-seq (scRNA-seq) dataset first published by Neftel et al. (2019).^8^ This dataset contains 5,689 single cells (4,865 neoplastic) across 20 adult GBM tumors. Neoplastic cells per tumor ranged from 121 to 435 cells. We applied a Module Score (MS) calculation to simultaneously score the entire GDS as a gene set in each single-cell in the Neftel dataset.^29^ We found that neoplastic GBM cells more highly express GDS genes than non-neoplastic tumor microenvironment (TME) cells in the Neftel dataset (Student’s t test, p < 0.0001, Figure 1D; non-neoplastic TME cells in this dataset include oligodendrocytes, T-cells, and macrophages).

To validate this finding, we performed the same MS calculations and comparisons on two additional scRNA-seq datasets of patient-derived GBM tumors – one published by Johnson et al. (2021) (28,130 cells, 5 adult IDH-wt GBM tumors)^30^ and one that we generated ourselves, in-house (42,839 cells, 6 adult GBM tumors). In each independent dataset, we similarly found that GDS genes were significantly more highly expressed in neoplastic compared to non-neoplastic single cells (Figures S3-S4). Moreover, to compare GDS expression specificity to GBM compared to other central nervous system (CNS) neoplasms, we analyzed a scRNA-seq dataset from Jackson et al. (2025) that contains single-cells from normal brain and patients with anaplastic astrocytoma, astrocytoma, oligodendroglioma, and GBM.^31^ Comparing normalized, tumor-level MS in cancer cells between CNS neoplasms emphasized the specificity of the GDS to GBM, with GBM tumors more highly expressing the GDS compared to anaplastic astrocytoma (ANOVA with Tukey’s post-hoc p < 0.05) and oligodendroglioma (ANOVA with Tukey’s post-hoc p < 0.01).

We then analyzed the pathways best represented in the 168-gene GDS. We performed pathway enrichment analysis using collections H, C2, and C3 of the Molecular Signatures Database (MSigDB) and found the GDS to be significantly enriched for pathways related to transcription and translation, adhesion, motility, and invasion (Figure 1E).^32–40^ Of the top 25 enriched terms among these collections, eight (8) were related to transcription and the translation machinery (e.g. “rRNA Processing (Reactome)”, “RNA Pol I Transcription Initiation [Termination] (Reactome)”). Additionally, 7 of 25 are related to mitosis and cell division (e.g. “Mitotic Metaphase & Anaphase (Reactome)”, “Nuclear Envelope Reassembly (Reactome)”, “Recruitment of Centrosome Complexes (Reactome)”), and 5 out of 25 are related to actin filaments and migration/adhesion (e.g. “Focal Adhesion (KEGG)”, “Folding of Actin by CCT/TriC (Reactome)”, and “RHO GTPases Activate WASPs & WAVEs (Reactome)”). Additional analysis of the GDS using the STRING database resulted in 4 clusters of interactions that corroborated pathway enrichments.^41^ Clusters included GDS genes enriched for rRNA metabolism, cell division, RHO GTPase activity, and COPII vesicle coating. Only 14 of 68 genes did not have any documented interactions with the other GDS genes (Figure S1E).

Therefore, the 168-gene GDS is not only specific to GBM cell lines but is more highly expressed in patient GBM tumor cells compared to non-neoplastic TME cells in those same tumors. The GDS is also more highly expressed in GBM compared to other CNS neoplasms. Moreover, these GDS genes are largely related to cellular mitosis, protein translation, and actin polymerization – key pathways in tumor development and progression.

### Glioblastoma dependency signature is required for identification of vulnerability states

We next sought to use the GDS to group single-cells by their patterns of expression of these genes. We hypothesized that cells clustered in such a way would share actionable vulnerabilities. For this analysis, we chose to again use the Neftel scRNA-seq dataset as it contains deep sequencing coverage on the largest number of patient tumors. The pipeline used for single-cell state identification is outlined in Figure 2A. For each of the 20 adult GBM tumors, we calculated the Jaccard similarity index (JS) for the binarized scaled expression of each GDS gene in every neoplastic cell. This initial step was performed on each tumor separately to minimize tumor-level effects and account for intertumor heterogeneity. Following Jaccard scoring, we performed hierarchical clustering on the resulting JS matrix for every cell in the tumor for all k from 2 through 30. A representative heatmap of this first step from tumor MGH106 is shown in Figure 2B. The remaining 19 JS heatmaps for each tumor are available in File S1.

**Figure 2:**
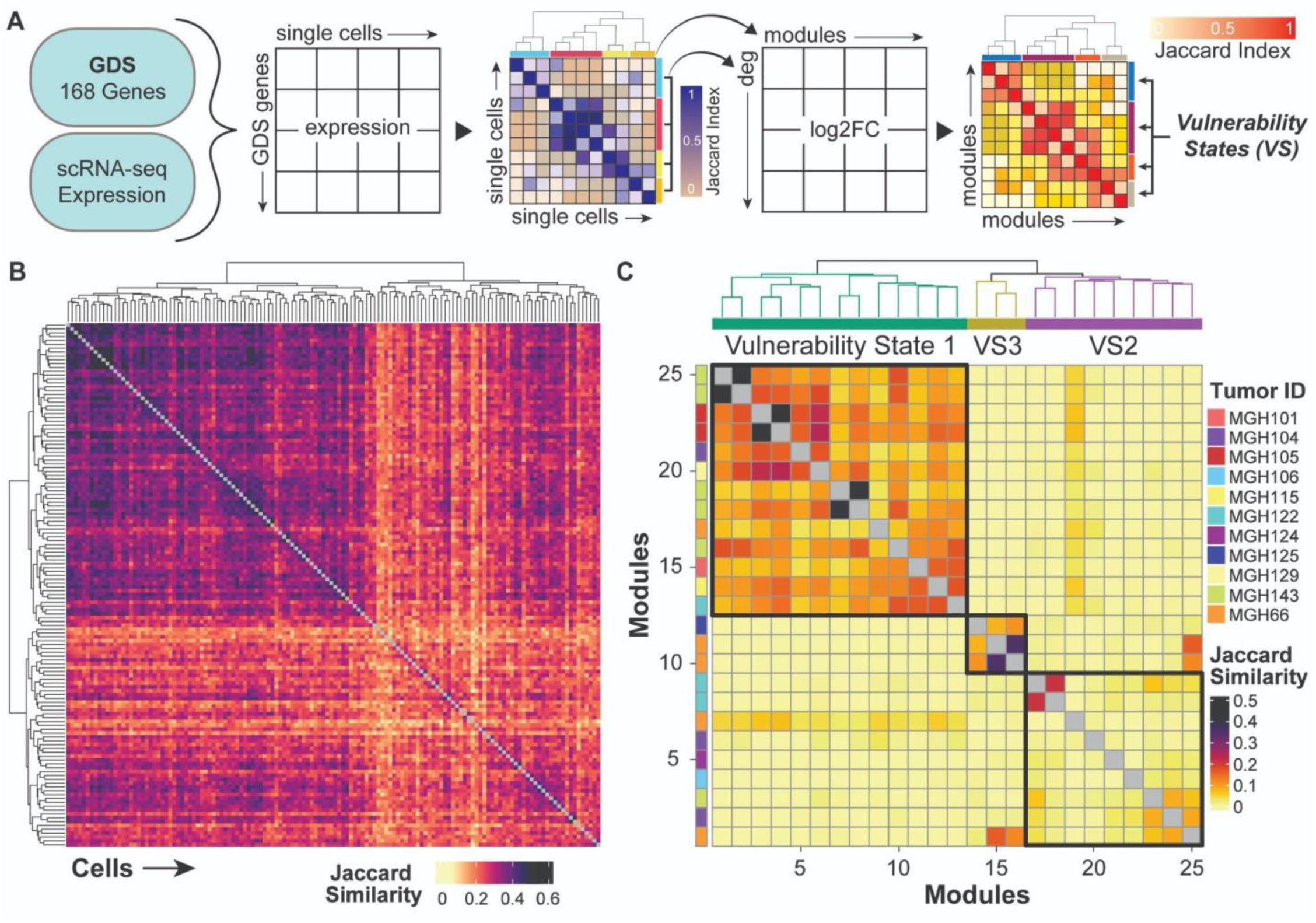
Glioblastoma dependency signature is required for identification of vulnerability states. (**A**) Schematic outlining the process of vulnerability state (VS) identification from single-cell expression data of glioblastoma dependency signature (GDS) genes (DEG = differentially expressed genes). (**B**) Representative heatmap of expression-derived Jaccard Index scores (Jaccard Similarity) for every neoplastic single-cell in one tumor (MGH106) out of 20 total adult GBM tumors in the Neftel et al. scRNA-seq dataset. Clustering of Jaccard scores was performed by Ward D2 hierarchical clustering at all k between 2 and 30 for this and all other tumors (File S1). Each row and column of the heatmap represents a single neoplastic cell in MGH106. (**C**) Heatmap representing Jaccard scores between pass-filter GDS-derived clusters (modules) across all 20 tumors. Pass-filter clusters included 25 modules from 11 distinct tumors. Each row and column of the heatmap represents one GDS-derived module from one tumor. The tumor identifier (Tumor ID) is used as row annotations to highlight which tumor each module is derived from. Clustering of Jaccard similarities between modules was performed by Ward D2 hierarchical clustering, identifying 3 final signatures (vulnerability states, VS).

We identified 274 clusters of single-cells across all 20 tumors with sufficient numbers of cells for downstream analysis. We then sought to collapse clusters based on shared transcriptomic profiles to account for overlap across k-values. To do so, we performed differential expression analysis between each cluster within a given tumor, using the top differentially expressed genes (DEG) with p < 0.05 and log2FC > 0 as a transcriptomic signature (module) for each cluster (*FindAllMarkers* from *Seurat* with MAST statistical testing).^29,42^ At minimum, a given cluster had to have 50 DEG that met these criteria, in addition to at least 1 gene with p < 0.001 to be used as a module. We then binarized modules based on whether a DEG was present or absent to then perform JS calculation to score each cluster for similarity to each other. We then removed any pairs of modules within a tumor that had JS > 0.75, retaining one of the two modules. This resulted in 25 unique modules across 11 (out of 20) tumors with unique GDS-informed transcriptomic signatures. Following this exclusion, we performed hierarchical clustering to collapse the remaining 25 modules into meta-clusters, which we termed vulnerability states (VS) (Figure 2C). We found 3 VS by this method. Clustering robustness and validity was confirmed via *clustree* S3 stability analysis (Figure S5A) and *clustree*-overlayed principal component analysis (PCA, Figure S5B). In conjunction with these visualizations, the optimal k-value (k = 3) was determined by calculation of the Calinski-Harabasz metric (Figure S5C). Importantly, rather than clustering by tumor identity, intertumor clusters group according to transcriptomic profiles (Figure 2C).

We next sought to identify marker genes for these cell states to allow for further characterization as well as cell state assignment and validation in additional data sets. The top 100 genes by log2FC across all modules within a given VS were selected as VS signature genes. Figure 3A shows the pseudo-bulk normalized scaled expression of the 292 unique identified signature genes across 24 of 25 modules from which they were derived. Differential enrichment analysis was then performed on these signature genes using the Molecular Signatures Database (MSigDB) collections H, C2, and C3 (Figure 3.3B).^32–40^ The results of this analysis accurately reflected the signature genes for each VS, with the VS1 signature enriching strongly for neural and developmental transcription factor (TF) gene sets, VS2 for transcription and translational machinery, and VS3 for cellular proliferation. Key VS signature genes that corroborate these enrichment results are highlighted in Figure 3A.

**Figure 3:**
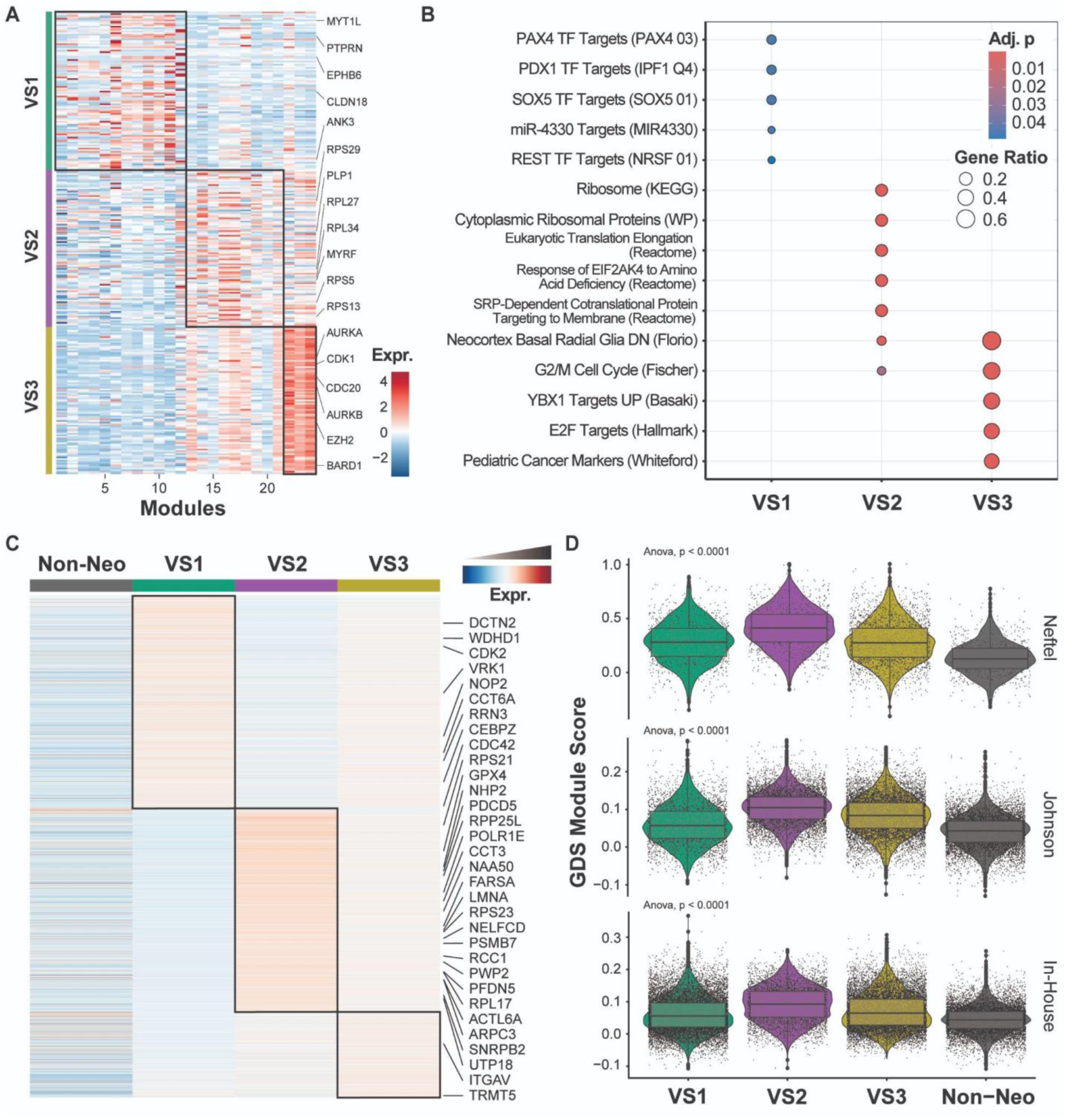
Glioblastoma vulnerability states are distinct. (**A**) Heatmap representing average scaled RNA expression of the n = 100 marker genes identified for each VS. Genes are ranked by descending log2FC for each VS. Labeled genes are those of interest with particular relationships to over-enriched terms identified in (B). (**B**) Dot plot displaying MSigDB differential enrichment analysis results (collections H, C2, and C3) performed on the 100 marker genes per VS as highlighted in (A) (n = 292 unique genes). The database each term is derived from within each collection is written in parentheses following the term name. Color indicates the BH-adjusted p-value and circle size represents the gene ratio. (**C**) The top 1,500 differentially expressed genes by log2 fold change (log2FC) between VS are visualized using a heatmap of scaled, normalized pseudo-bulk gene expression. Total DEG identified for VS3 was 634. Labeled genes are GDS genes (32 out of 168) that are also significantly differentially expressed. DEG were identified using Seurat and the MAST statistical test with a Bonferroni-adjusted p-value < 0.05 and log2FC > 0.0. (**D**) Violin plots displaying MS of the 168-gene GDS in single-cells in each of the datasets used in the study (Neftel, Johnson, and Suter, respectively), with cells grouped by their assigned VS. Each black dot represents one single-cell in the dataset. Statistical testing was performed by ANOVA with Tukey’s post-hoc test for individual comparisons and presented as a Bonferroni-adjusted p-value.

To identify DEG between each VS and further characterize these novel cell states, we then performed differential expression analysis. Differential expression profiles between each VS were distinct, with more GDS genes identified as DEG in VS2 than either VS1 or VS3 others (Figure 3C). In total, 32 of 168 GDS genes were identified among the top 1,500 significant DEG for each VS (p < 0.05, log2FC > 0, genes annotated in Figure 3C). VS1 differentially expressed 9 GDS genes: DCTN2, WDHD1, CDK2, VRK1, NOP2, CCT6A, RRN3, CEBPZ, and CDC42. VS2 differentially expressed 21 GDS genes including LMNA and RCC1. VS3 differentially expressed 2 GDS genes: ITGAV and TRMT5. Notably, VS2 had the largest number of GDS genes represented among its significant DEG.

Moreover, Gene Ontology (MSigDB collections H and C5) differential enrichment analysis performed on the top 100 significant DEG (p < 0.05) with log2FC > 0.5 (Figure S6), revealed that each VS had distinctly enriched pathways that mirrored VS signature gene enrichment (Figure 3B). VS2 was more highly enriched for pathways related to extracellular matrix organization. By contrast, VS1 DEGs are enriched for apoptotic and trans-synaptic signaling regulation in addition to transcriptional activator activity, while VS3 DEGs are related to exocytotic pathways (e.g. synaptic exocytosis).

We are therefore able to use GDS genes to inform identification of GBM VS. Moreover, GDS genes are both present and reflected in both the VS signature genes and the significant DEG between VS, especially when comparing our analyses demonstrated in Figure 1D with those in Figures 3B-C, S1, & S6. Taken together, VS have distinctive signatures and transcriptional profiles. VS1 largely reflects a stem-like transcriptional signature, VS2 is related to translational machinery and largely recapitulates the GDS (Figure 3D), and VS3 is related to the cell cycle and exocytosis (e.g. platelet alpha granule GOCC).

### Vulnerability states are present across multiple scRNA-seq datasets

We next sought to validate the presence of VS in additional GBM tumors and to further study their single-cell transcriptomic profiles. To that end we downloaded, processed, and analyzed a GBM scRNA-seq dataset from Johnson et al. to use in conjunction with our own in-house dataset. The Johnson dataset contains 28,130 single cells (19,530 neoplastic) from 5 adult GBM tumors (Figure S3).^30^ Our in-house dataset contains 42,839 single cells (32,159 neoplastic) from 6 adult GBM tumors (Figure S4).

We used VS signature genes (Table S4) to score single-cells in each dataset for enrichment of those genes using *singscore*, an independent-sample gene-set scoring method.^43,44^ *Singscore* enrichment scores were then used for cell state assignment, with cells being assigned to whichever VS scored most highly in that cell. VS were identified in both datasets, and the resulting VS proportions by tumor in both the Johnson and our in-house dataset largely recapitulated the proportions observed in the Neftel dataset. Individual tumors in all three datasets differed slightly in the relative proportions of their VS content, with some more highly or more lowly containing some VS than others. As a whole, VS3 makes up the majority of the neoplastic cell populations, followed by VS1, and then by VS2 (Figure 4A).

**Figure 4:**
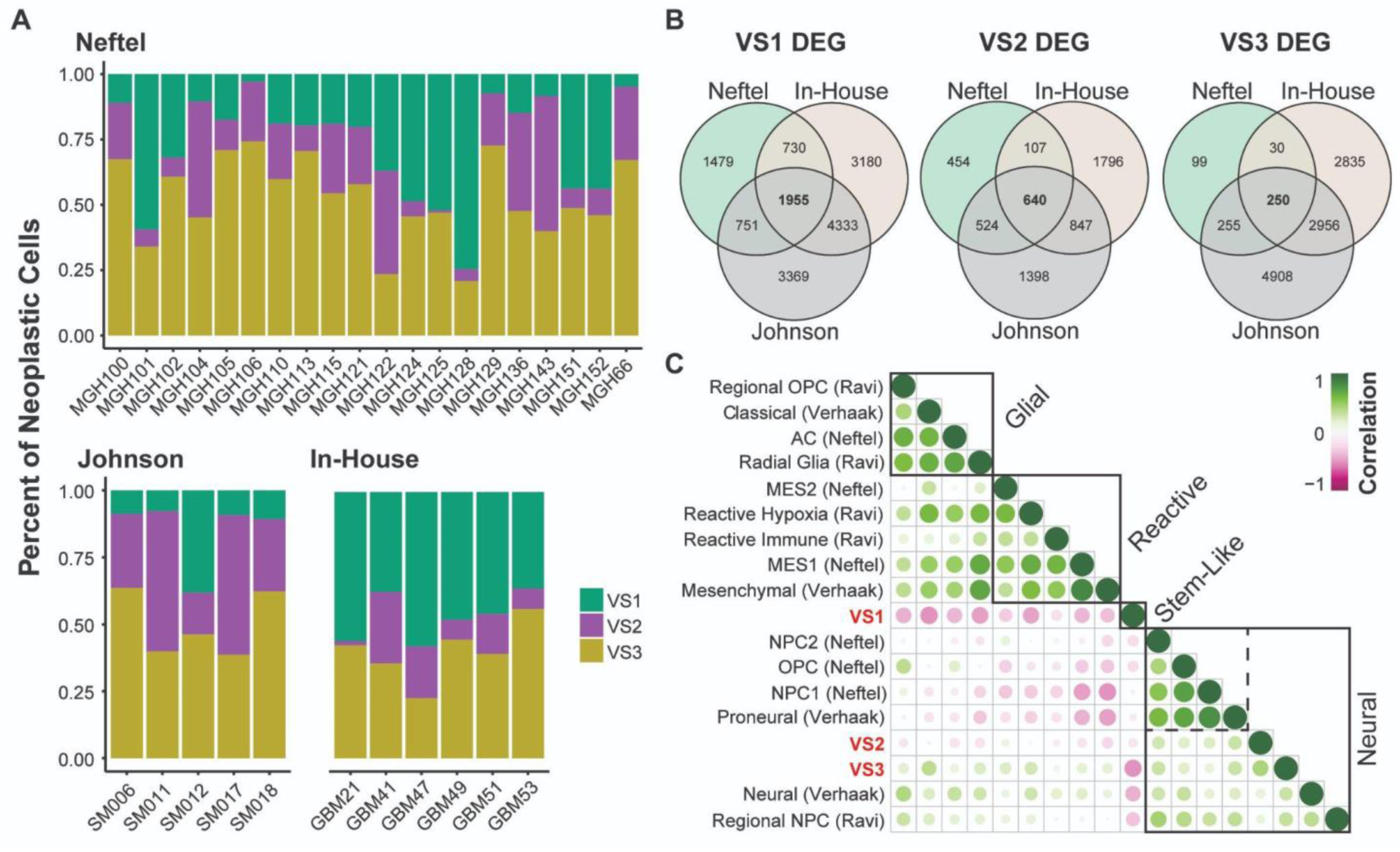
Vulnerability states are present across multiple scRNA-seq datasets. (**A**) Stacked bar plots displaying the proportions of neoplastic cells that are categorized as each VS for each tumor in the Neftel et al., Johnson et al., and our in-house scRNA-seq datasets, respectively. Color corresponds to VS. VS assignments in external datasets were made using singscore enrichment scoring. (**B**) Venn diagrams representing overlapping DEG among each of the 3 scRNA-seq represented in A. For each dataset, DEG were identified between VS using Seurat’s FindMarkers function with log2FC cutoff of 0 and significant p-value < 0.05 as determined by MAST statistical testing. The number of DEG in each category are represented here with the percentage of total DEG per VS among all 3 datasets shown in parentheses. (**C**) Spearman correlation analysis of singscore enrichment scores between existing GBM transcriptional signatures (from Verhaak et al. 2010; Neftel et al. 2019; and Ravi et al. 2022) and our own vulnerability state signatures (red). Clustering of all signatures produces 4 major groups (with 1 sub-group): Glial-like, Reactive-like, Stem-like (VS1), and Neural-like (including VS2 and VS3 within their own subcluster). Enrichment scores calculated from our In-House scRNA-seq dataset.

Given the consistency of VS scoring and assignment in each dataset, we wanted to validate the similarities between assigned VS across datasets, independent of the 100-gene VS signatures. To do so, we performed differential expression analysis between VS in the Johnson and in-house datasets, respectively, with the aim of comparing the resulting DEG with those identified in the Neftel dataset. DEG were calculated using *Seurat’s FindAllMarkers* function and determined by MAST-adjusted p-value < 0.05 and log2FC > 0. For VS1 across all three scRNA-seq datasets, a total of 1,955 (12.4% of all VS1 DEG) of DEG overlapped. Similarly, for VS2 a total of 640 (11.1%) of DEG overlapped. For VS3, 250 genes overlapped though this only represented 2.21% of VS3 DEG across all three datasets. However, the VS3 DEG identified in the Johnson and in-house dataset had high concordance, with 3,206 (28.3%) overlapping genes. These findings emphasize that the VS-assigned cells identified in the Johnson and our in-house datasets are similar to each other and to VS-assigned cells in the Neftel dataset, allowing us to conclude that VS are not a phenomenon isolated to the dataset from which they were derived.

Once confident in our validation of VS assignments, we finally sought to return to the GDS; since the premise of our investigation was to identify cell states with shared GDS expression patterns, we wanted to identify whether VS express GDS genes in all datasets. We used the GDS MS calculated previously for each single cell in all three datasets and grouped cells by neoplastic identity, comparing GDS expression between assigned VS. As highlighted in Figure 3C and quantified in Figure 3D, VS differently express GDS genes, with VS2 more highly expressing GDS genes compared to both VS1 and VS3 in all 3 datasets (ANOVA p < 0.0001 for all three datasets; VS2 MS > VS1 and VS3 MS by Tukey’s post-hoc test, p < 0.0001 for all three datasets; Figure 3D).

Transcriptional signatures representing bulk, single-cell, and spatial transcriptomic classification systems for GBM from Verhaak et al., Neftel et al., and Ravi et al., respectively, were acquired with the goal of comparing these to our VS signatures.^8,15,45^ Each signature was scored for enrichment in every neoplastic single-cell in our in-house dataset using *singscore*. Enrichment scores were compared using Spearman correlation to identify similarities between signatures (Figure 4C). Our results largely recapitulated the results observed in Ravi et al. (2022).^15^ However, the localization and placement of our VS signatures in this plot are critical. First, we observe similar grouping of glial and reactive (immune response, hypoxia) signatures. While Ravi et al. (2022) observed clustering of one neural group, we see a larger neural cluster with a sub-cluster (dotted lines) owing to our identification and use of VS2 and VS3 signatures. While neural-like signatures by this analysis, VS2 and VS3 are unique enough to be distinct from Neftel NPC and OPC states and Verhaak Proneural tumor type, yet similar to the regional neural progenitor-like state observed in the spatial transcriptomic analysis conducted by Ravi et al. (2022). Critically, VS1 was not similar to any of the existing signatures, as it clustered independently with low Spearman correlations to all other transcriptional signatures. Importantly, the overlapping DEG from Figure 4B enriched for pathways that corroborate this dissimilarity (Figure 6D-F, Figure S7A-C). Based on VS signature gene enrichment analysis (Figures 3A-B), VS DEG enrichment analysis (Figure 6A & 6D-F, Figures S6-7), and our findings here (Figure 4C), VS1 most likely represents a novel neural stem-like signature in GBM.

These findings emphasize that VS are found across GBM tumors, but precise proportions of these cell states differ between patients. Despite this proportional variance, their concordance across tumors is high. Moreover, VS2 shows the highest degree of similarity to the GDS signature. By comparing transcriptional classification signatures from previously-published literature, we show that VS1 is a unique stem-like classification signature for cell states in GBM. We also find that VS2 and VS3 represent neural-like signatures related to transcription/translation and exocytosis and the cell cycle, respectively.

### Glioblastoma vulnerability states are spatially exclusive

Given our findings using GBM transcriptomic classification signatures from the literature, particularly those from Ravi et al. and a growing body of research emphasizing the importance of cellular spatial distribution and heterogeneity in GBM, we sought to investigate how our identified VS were distributed spatially in patient tumors.^12,15,46,47^ Corroborating our data in spatial samples is essential given continuously emerging evidence that spatial cellular organization plays a role in driving clonal evolution, tumor progression, and treatment resistance.

We obtained the publicly-available spatial transcriptomic (stRNA-seq) data published by Ravi et al. through Dryad (“Dryad” 2025) and processed and analyzed raw counts according to our in-house protocols. These data contained 18 stRNA-seq GBM tumor sections. Our in-house scRNA-seq dataset was then used to deconvolute the bulk RNA-seq spots of the 10X Visium stRNA-seq data using *SCDC* to glean insight into the spatial distribution of vulnerability states in these tumor slices.^48^ Our deconvolution results highlight that spatial distribution of VS varies from tumor-to-tumor, with most tumors having lower levels of VS1 present while being either VS2-high or VS3-high, relatively. These findings are quantified in Figure 5A with deconvoluted proportions of non-neoplastic cells and VS presented across all 18 tumor sections. The SCDC deconvoluted VS proportions for all 18 tumor sections are provided in Figures S9.

**Figure 5:**
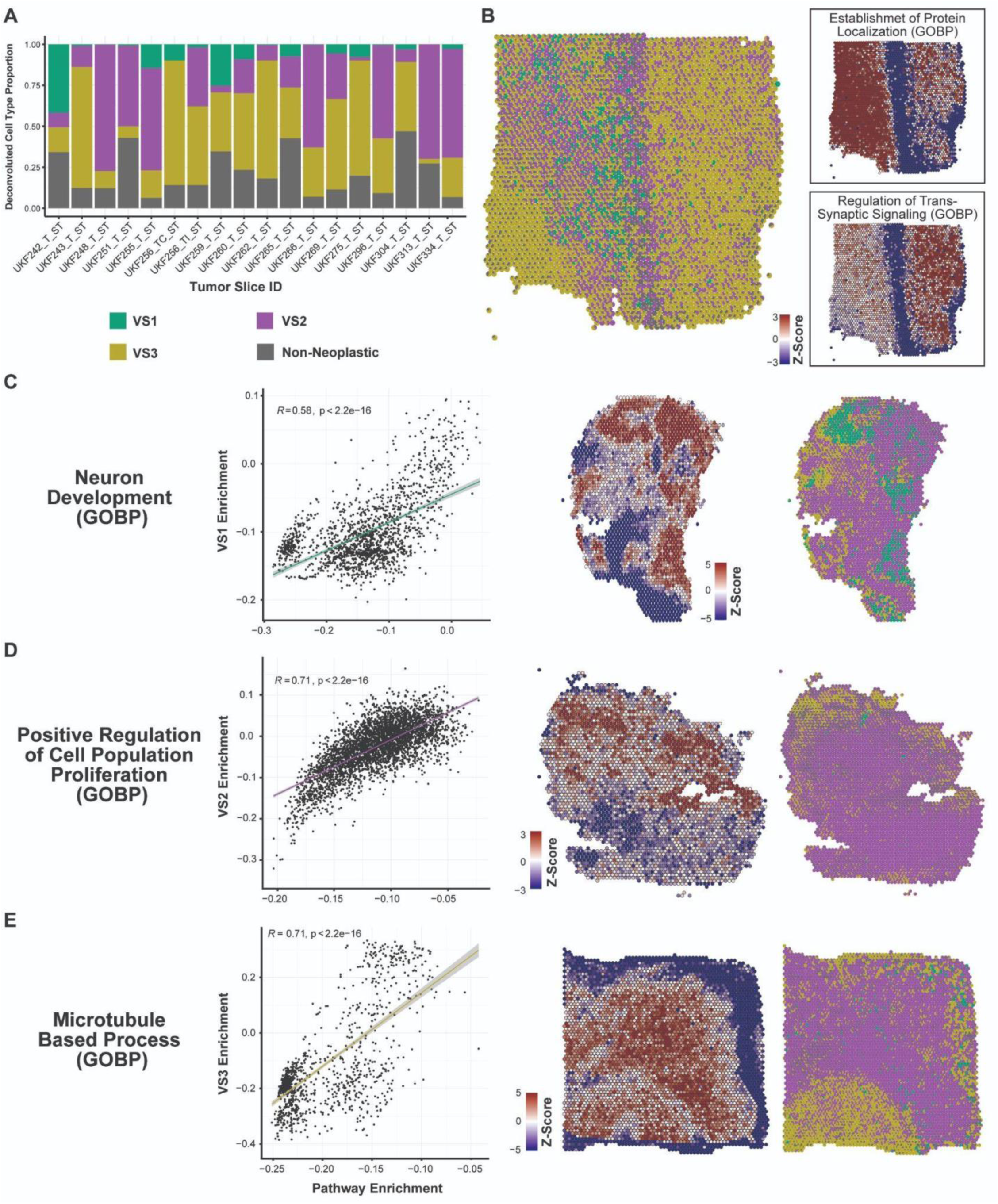
Glioblastoma vulnerability states are spatially exclusive. (**A**) 10X Visium stRNA-seq data were acquired from Ravi et al. and deconvoluted using SCDC to identify single-cell type proportions present in each section, with our in-house scRNA-seq dataset as a reference. This stacked bar plot shows the deconvoluted cell type proportions present within each of 18 GBM tumor sections. Cell types included as “Non-Neoplastic” designation include: astrocytes, endothelial cells, fibroblasts, myeloid cells, oligodendrocytes, and T-cells. (**B**) Shows a spatial scatter plot of UKF269 where each spot is represented as a pie chart showing the proportion of cell types identified by deconvolution at that precise location. To the right of the spatial scatter-pie image are two representative images of gene set co-regulation analysis (GESECA), with respective enrichment scores at each spot represented as z-scores. (**C-E**) Display, in succession, select top-upregulated pathways for each of VS1 (C), VS2 (D), and VS3 (E), visualized by spot-level enrichment score correlations, GESECA z-score spatial scatter plots, and spatial scatter-pie images, from left to right. (C) specifically shows VS1 enrichment for Neuron Development, (D) shows VS2 enrichment for Regulation of Cell Population Proliferation, and (E) shows VS3 enrichment for Microtubule Based Processes. All enrichments were performed using *fgsea* using Gene Ontology Biological Process (GOBP) enrichment set, with correlation analysis performed using Pearson correlation with a level of significance set to < 0.05.

In addition to tumor sections varying by proportion of VS present, spatial distributions of Gene Ontology Biological Processes (GOBP) enriched pathways were unique and overlapped with VS distributions. We performed gene set co-regulation analysis (GESECA) implemented through the *fgsea* R package on each tumor section independently using the GOBP database.^49^ The goal was to identify spatial patterns of gene set expression. For example, we found that one section (UKF269) showed a stark contrast between cells involved in protein localization (Establishment of Protein Localization) compared to those involved in trans-synaptic signaling (Regulation of Trans-Synaptic Signaling) (Figure 5B). Such contrasts are also interesting when examining relative VS proportions in each of these enriched regions – VS1 and VS2 are more present in tissue spaces involved in protein localization while VS3 appears more involved in trans-synaptic signaling unless specifically sharing space with non-neoplastic cell types.

We next sought to correlate spot-level VS enrichment to pathway enrichment scores in a spatial landscape. We took the top regulated pathways in each section from our GESECA analysis, scored every spot for enrichment of those pathways, and concurrently scored each spot for VS enrichment using *singscore*.^43,44^ We then ran correlation analyses using Pearson’s R to compare VS enrichment with GOBP pathway enrichment directly, across all spots among all sections. Among these comparisons, those pathways with significant correlations to VS enrichment were strikingly similar to our previous scRNA-seq-based enrichment results. VS1 strongly correlated with GOBP terms related to cellular and neural lineage development and differentiation (“Neuron Development”, R = 0.58, p < 0.0001, Figure 5C; “Neuron Differentiation”, R = 0.54, p < 0.0001, Figure S8). Additional enrichments were for synaptic transmission and synaptic plasticity (“Inhibitory Synapse Assembly”, R = 0.83, p < 0.0001, Figure S8). VS2 strongly correlated with terms related to cellular proliferation, response to cellular stress, and regulation of cell death, reinforcing our findings that VS2 most highly expressed GDS genes (“Positive Regulation of Cell Population Proliferation”, R = 0.71, p < 0.0001, Figure 5D; “Cellular Response to Stress”, R = 0.73, p < 0.0001; “Regulation of Cell Death”, R = 0.8, p < 0.0001, Figure S8). Interestingly, VS2 enrichment also correlated with immune and inflammatory response-related pathways. Lastly, VS3 enrichment correlated with microtubule-related pathways (“Microtubule Based Process”, R = 0.71, p < 0.0001, Figure 5E; “Transport Along Microtubule”, R = 0.69, p < 0.0001, Figure S8) and pathways related to oxidative phosphorylation including cellular aerobic respiration and ATP production (“Aerobic Respiration”, R = 0.79, p < 0.0001, Figure S8).

Importantly, we observed that expression of two genes our laboratory has recently reported to be important in GBM metabolism (MAT2A and AHCY, Rowland et al. 2025) are spatially localized to VS2 (Figures S9-11), which is in line with our findings that VS2 contains genes related to metabolic pathways. DepMap data suggested that MAT2A and AHCY knockdown in GBM cell lines reduced survival. We extended these findings to more relevant GBM PDX lines and glioma stem cells, suggesting that MAT2A and AHCY are essential genes in GBM and that our overall approach to DepMap-derived GBM dependencies is robust.

These results strongly corroborate our findings from analysis of the scRNA-seq data – that VS distinctly express transcriptional programs important to GBM – while revealing their spatial correlations with critical tumorigenic pathways. Additionally, the proportions of VS present in a given tumor differ in all tumors analyzed by both scRNA-and stRNA-seq. This could be leveraged for patient stratification pending deeper interrogation of VS functional and biological significance, particularly with regard to response to therapy.

### Vulnerability states express differently actionable transcriptional programs

Our analysis of VS across multiple scRNA-seq datasets in addition to our findings from spatial analysis spurred deeper investigation to understand the therapeutic implications of VS. We performed differential enrichment analysis of the top 300 DEG between VS in the Neftel data using the KEGG Medicus database and found VS1 to be significantly enriched for cell cycle and replication pathways (e.g. Homologous Recombination, Origin Unwinding and Elongation), while VS2 enriched for translation and metabolism terms (e.g. Translation Initiation, Electron Transfer in Complex I), and VS3 enriched for pathways related to cytoskeleton assembly, migration, and adhesion (e.g. ITGAB/FAK/CDC42 Signaling, ITGAB/RHOG/RAC Signaling) (Figure 6A).

**Figure 6.**
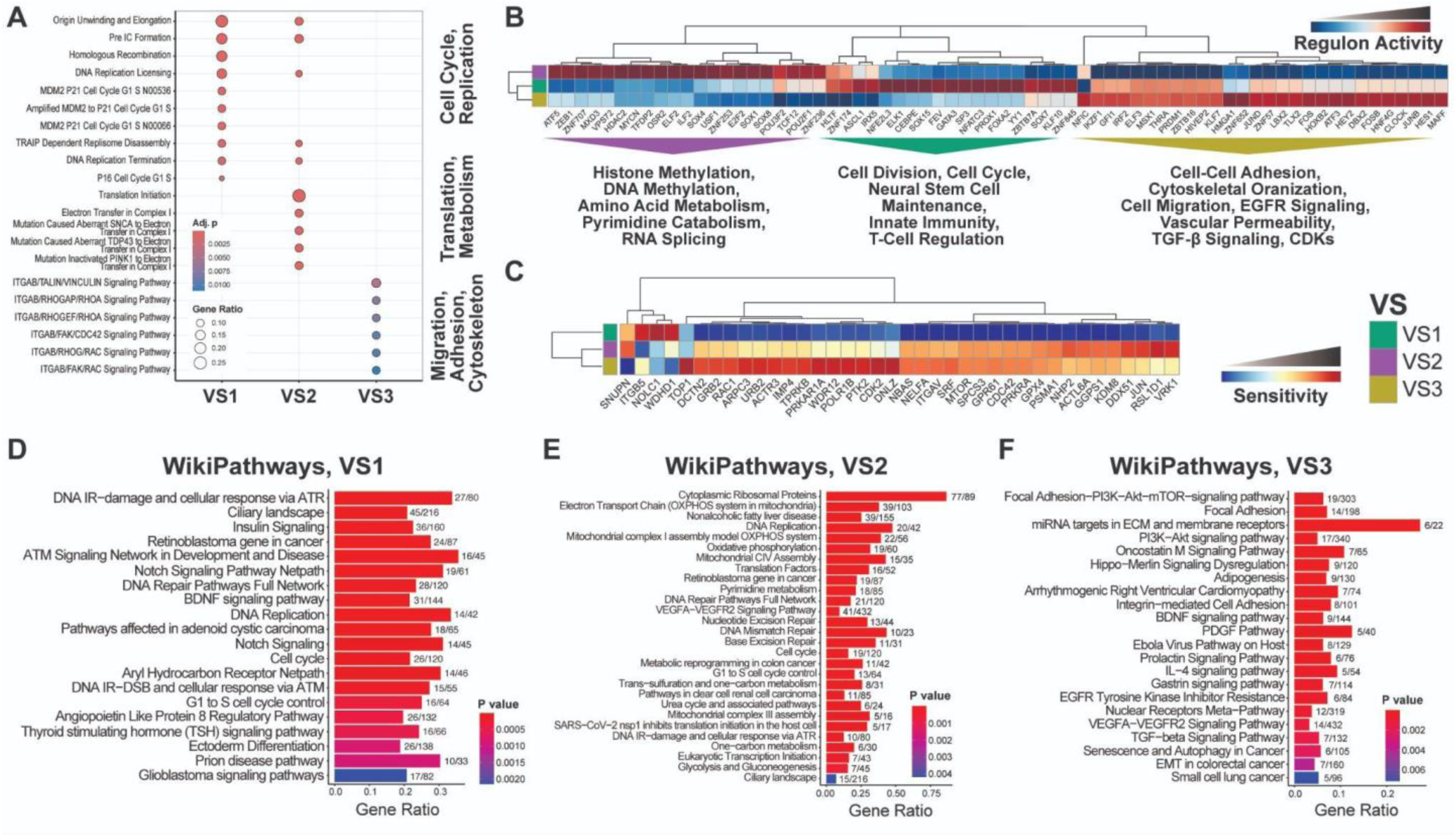
Vulnerability states express differently actionable transcriptional programs. (**A**) Dot plot showing the results of KEGG Medicus differential enrichment analysis performed on the top 300 differentially expressed genes for each VS. Color indicates the BH-adjusted p-value and circle size represents the gene ratio. (**B**) Heatmap displaying the relative activity scores of the top 25 significantly different regulons for each VS by SCENIC analysis. (**C**) Predicted sensitivities of each VS to targeted GDS CRISPR-mediated knockout (38 / 168 GDS genes analyzed). (**D-F**) Bar plot of enrichment results for all overlapping differentially expressed genes for VS1, VS2, and VS3 (respectively) among the Neftel et al. (2019), Johnson et al. (2020), and our in-house scRNA-seq datasets. Analysis conducted using the WikiPathways gene set. Color indicates BH-adjusted p-value by Fisher’s exact test with gene ratio plotted on the x-axis (fractional representation to the right of each bar).

Subsequent investigation into the transcriptional drivers of these three cell states using SCENIC corroborated these findings (Figure 6B).^50^ Increased activity of transcription factors (TFs) such as JUND, FOS, IKZF1, THRA, and KLF7 emphasized the migration, adhesion, and cytoskeletal functions identified in VS3. Moreover, MYCN, ELF2, SOX1, and SOX4 activity in VS2 highlight the metabolic roles that these cells play. Lastly, SP3, FOXA2, and ASCL1 activities in VS1 cells show that these cells are involved in cell cycle signaling, neural stem cell maintenance, and interactions with the immune system.

The LINCS L1000 dataset contains transcriptomic data from thousands of cell line perturbations, including small molecule inhibition, siRNA knockdowns, and CRISPR knockouts.^51–53^ We probed this dataset and found CRISPR knockout data for 39 of 168 GDS genes. For those 39 genes, we then leveraged the RNA-seq data of all cell lines perturbed by CRISPR knockout in the L1000 dataset to identify a transcriptional consensus signature (TCS) for each gene that comprises the aggregate top up-and down-regulated genes in response to perturbation as previously described.^54^ We identified TCSs for 38 of 39 genes, owing to insufficient cell lines perturbed by 1 gene. Each neoplastic sing-cell across all three scRNA-seq datasets were then scored for expression of the TCS of each gene knockout using *singscore*. Using the principle of disease signature reversal, those single-cells that scored most highly for a TCS were predicted to be resistant to that knockout, whereas those that score most lowly for a TCS are predicted to be sensitive to gene knockout. CRISPR perturbation scores (XPS) were then aggregated and normalized for each single-cell across datasets, and the pseudobulk average XPS for each gene was calculated for each cell state. By this analysis, we found that VS were differently sensitive to targeted GDS knockout (Figure 6C). Though VS2 most highly expresses GDS genes, it is only most sensitive to a handful of the 38 GDS genes, most notably JUN and VRK1. VS1 was found to only be sensitive to targeted knockout of ITGB5, NOLC1, and WDHD1. By contrast, VS3 had multiple notable sensitivities, including RAC1, TOP1, CDK2, PTK2. Both VS2 and VS3 are predicted to be moderately sensitive to MTOR and GPX4 knockout, though VS1 has predicted resistance to both. These data support previously published literature showing that monotherapy with RAC1 inhibitors, CDK inhibitors, and TOP1 inhibitors have inconsistent efficacy in GBM.^55–58^

We additionally performed enrichment for all overlapping DEG between scRNA-seq datasets presented in Figure 4B. We found that the enriched pathways by MSigDB analysis reflect actionable pathways including Wnt-Beta catenin signaling and Hedgehog signaling for VS1, MYC targets and MTOR signaling for VS2, and TGFB1 signaling for VS3 (Figure S7A-C). Deeper investigation of enriched pathways using the WikiPathways database and only protein-coding genes revealed overlap with the aforementioned enrichments, predicted XPS sensitivities, and regulon activities for each VS. Notable terms enriched in VS1 that also included GDS genes were “DNA Replication,” “Cell Cycle,” and “BDNF Signaling Pathway” (Figure 6D). For VS2, “Pyrimidine Metabolism” and “Eukaryotic Transcription Initiation” are notable (Figure 6E). For VS3, enriched terms included “Focal Adhesion” and “PI3K-AKT Signaling Pathway” (Figure 6F).

Taken together, this series of analyses reinforces that VS are differently actionable and that the trio of enrichments, CRISPR knockout sensitivities, and SCENIC-based regulon analysis produce similar candidate pathways for targeted therapy.

### Glioblastoma vulnerability states are differently sensitive to chemotherapy and small molecules

To test the LINCS L1000 and enrichment-derived functional predictions, we first leveraged ISOSCELES (Suter et al. 2025, *bioRxiv*) to identify whether approved small molecules for treatment of cancer patients aligned with these data and then tested select compounds and pathways using existing in vivo scRNA-seq data of treated GBM PDX.

ISOSCELES analysis revealed compounds such as palbociclib (CDK inhibitor) and etoposide (topoisomerase inhibitor) have the highest predicted sensitivities in VS3, in concordance with predicted sensitivities derived from analysis of the LINCS L1000 data (Figure 7A, Figure 6C). VS3 also displayed enhanced predicted sensitivities to known microtubule and migration inhibitors such as ixabepilone and cabazitaxel. Additionally, selective sensitivity of VS1 to inhibitors of DNA and RNA synthesis (e.g. capecitabine, ftorafur, and gimeracil) align with the presented enrichment and regulon activity results showing high cell cycle activities among VS1 cells. Importantly, VS2 showed selective sensitivity to TMZ with VS1 and VS3 predicted to be resistant.

**Figure 7.**
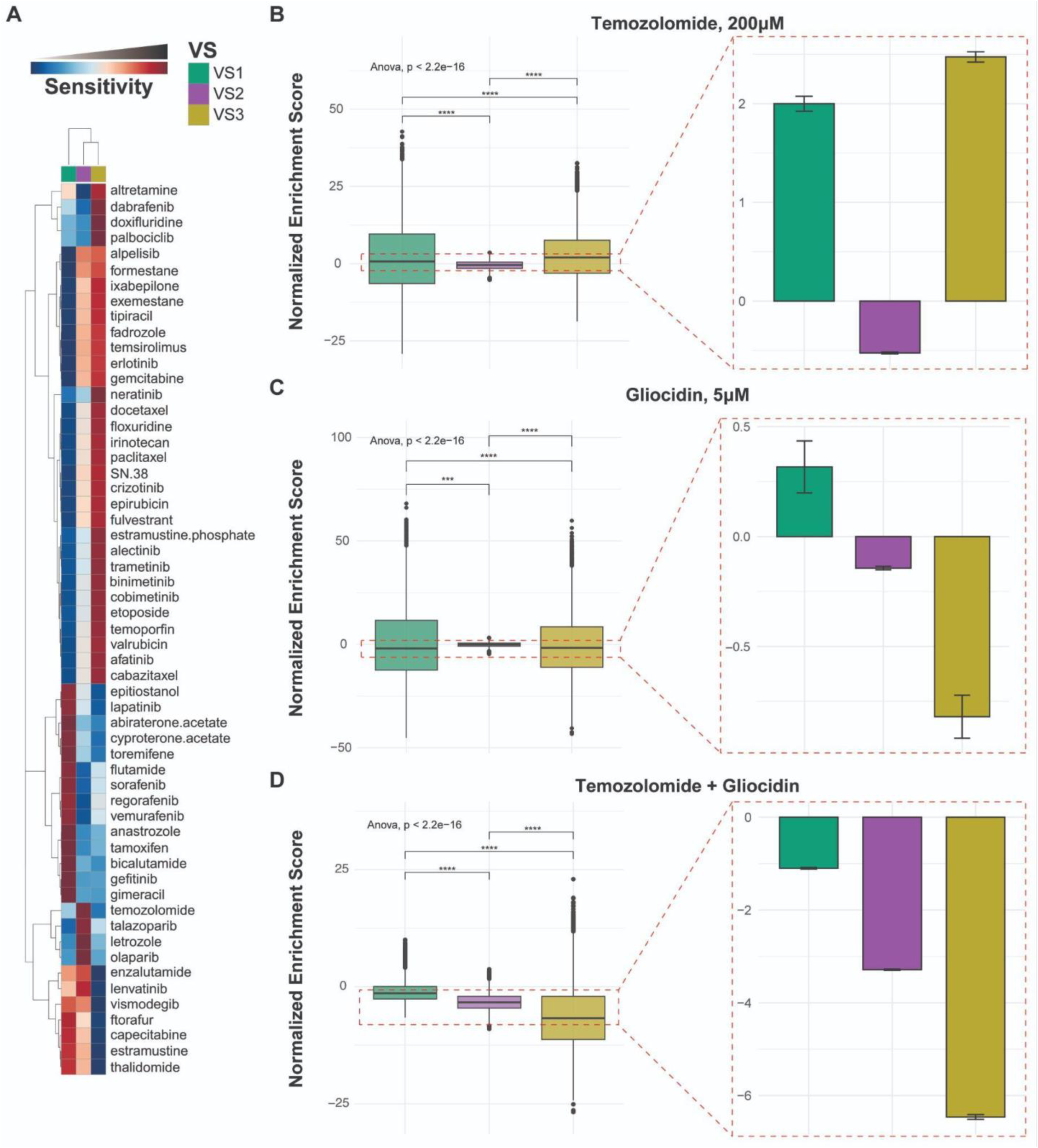
Glioblastoma vulnerability states are differently sensitive to chemotherapy and small molecules. (**A**) Heamap displaying predicted VS sensitivities to drugs FDA approved for use in oncology. B-D. Barplots showing single-cell VS enrichment scores for GBM PDX treated in vivo for 10 days with TMZ (**B**), gliocidin (**C**), or combination (**D**) at the noted dosages. Data indicate DMSO-and VS-proportion-normalized means +/-SD for n = 3 pooled biological replicates. Statistical analysis between VS enrichments was performed using ANOVA with Tukey’s post-hoc test (**** p < 0.0001). scRNA-seq data downloaded from Chen et al. (2024).

Given this information, we leveraged a recently-published scRNA-seq dataset of GBM PDX treated *in vivo* with TMZ, gliocidin (guanine synthesis inhibitor), and a combination of both to test our prediction that VS2 would be selectively targeted by TMZ and that gliocidin would target VS2 due to enrichments for nucleotide metabolism (Figure 7A, Figure 6E).^59^ The Chen et al. (2024) dataset contains scRNA-seq data from n = 3 biological replicates from each of four treatment groups: 200 µM TMZ, 5 µM gliocidin, 200 µM TMZ + 5 µM gliocidin, and DMSO. VS normalized enrichment scores for each treatment group revealed TMZ-treated tumors were significantly depleted in VS2 enrichment, and gliocidin-treated tumors were significantly depleted for both VS2 and VS3 enrichments (Figure 7B). Notably, the combination treatment displayed significant depletion of all 3 VS, emphasizing the importance of dual or combination therapy for effective GBM treatment. We additionally observed VS-specific compound sensitivities in both scRNA-seq and stRNA-seq data that support the *in vivo* synergy observed between CT-179 (OLIG2 inhibitor) and Depatux-M (Depatuxizumab Mafodotin, an antibody drug-conjugate consisting of an EGFR antibody conjugated to MMAF, a tubulin inhibitor) in Suter et al. (2025).^91^ Here, we observe CT-179 (VS1 targeting), microtubule inhibitors (docetaxel, paclitaxel; VS3 targeting), and EGFR inhibitors (afatinib, erlotinib; VS3 targeting) target distinct VS (Figure 7, Figures S12-S15).

## DISCUSSION

Transcriptional cell states and state transitions have been described as viable therapeutic targets due to their roles in drug resistance, tumor progression, immune evasion, and recurrence.^8,12–14,30,46,60–63^ However, it remains unclear which cells to target and prioritize. Here, we present a novel framework for cell state identification that is informed by lineage-specific vulnerabilities in GBM. Analysis of genome-wide cell line CRISPR screen data from the DepMap Consortium yielded a 168-gene dependency signature for GBM that we leveraged together with scRNA-seq data to identify shared GDS-related transcriptional modules in cells across patient tumors using similarity scoring and iterative hierarchical clustering. We demonstrate that the GDS is GBM-specific both compared to other cancer lineages and to other CNS neoplasms and that it is relevant to patient tumors in both scRNA-seq and stRNA-seq datasets. Moreover, we show that the GDS-derived VS are distinct from existing GBM tumor and cell type classification signatures in both scRNA-seq and stRNA-seq data and that they share spatial colocalization and enrichment with critical GBM tumorigenic pathways. Lastly, we demonstrate that VS have distinct biological functions with specific relevance to standard-of-care GBM treatment. Taken together, our results reveal that lineage-specific essential genes inform biologically significant and therapeutically relevant cell states in GBM.

DepMap and Project Achilles’ CRISPR screen data have been used to successfully identify candidate genetic targets for multiple cancer types.^25–28^ Gene essentiality is a measure used to quantify the effect of gene loss on a model organism or system. We hypothesized that we could use gene essentiality and single-cell expression in GBM to identify cells to prioritize for drug development. Using DepMap’s CRISPR data, we identified 168 genes specifically essential to GBM cell lines. Our GDS complements existing literature showing that some of our top GDS hits such as LMNA, JUN, RAC1, and CDC42 have key roles in GBM aggression and tumorigenicity.^64–67^ Translating *in vitro* results to patient data is challenging, yet our investigation highlighted GDS specificity to GBM neoplastic single-cells. Moreover, we identified minimal overlap between the GDS and dependency signatures from other tumor lineages and showed GDS specificity to GBM compared to other CNS neoplasms (Figure S1). Even with the inclusion of common essential genes, we would have expected greater overlaps between dependency signatures if lineage-specific gene essentiality were merely an *in vitro* phenomenon.

Transcriptional ITH in GBM has been explored extensively, with researchers characterizing cell states in high resolution using both scRNA- and stRNA-seq technologies.^8,9,12–15,30,63^ We show that our identified VS correlate with – yet are distinct from – some of the leading GBM transcriptomic classification signatures. Our stem-like signature, VS1, critically, does not positively correlate with any modules from Neftel et al., nor those from Ravi et al.’s spatial transcriptomic analysis or Verhaak et al.’s bulk RNA-seq-based GBM subtype classification scheme.^8,9,45^ It does, however, exhibit strong neural enrichment compounded with upregulation of canonical stem-cell pathways and embryonic TFs (PAX4, SOX5, SOX7, FOXA2, REST), including the Wnt/Beta-catenin and Hedgehog signaling pathways. Therefore, VS1 most likely represents a unique population of neoplastic, neural stem cell-like cells despite it not clustering with the NPC1 or NPC2 signatures from Neftel et al. Drexler et al. recently described a high-neural epigenetic signature in GBM that was correlated with poorer patient outcome, higher glioma-neural synaptic integration, and high levels of neural stem cell-like cells, which may be a starting point toward exploring the functional significance of our VS1 cell signature for exploitation toward drug development.^68^ Already, our analysis demonstrates predicted sensitivity to DNA and RNA synthesis inhibitors.

By contrast to VS1, our VS2 and VS3 signatures do correlate with cell state classifications from Ravi et al. (Regional NPC) and Neftel et al. (NPC1, NPC2, OPC) in addition to Verhaak et al.’s Proneural and Neural GBM transcriptional subtypes. In their publication Neftel et al. largely attribute NPC-like states to CDK4 amplification and OPC-like states to PDGFRA amplification, and subsequent research largely corroborates this.^14,15^ VS2 signatures are associated with translation and metabolism in GBM, while VS3 signatures are associated with migration, adhesion, and cytoskeletal pathways. We additionally show, however, that VS2 is strongly and uniquely GDS-like, expressing the most GDS-derived genes and sharing overlapping enrichment profiles with the GDS. Given its additional spatial-level correlations with anti-apoptotic and immunoregulatory biological processes and its selective sensitivity to TMZ, VS2, while sharing qualities of NPC-like cells, is distinct from them. Moreover, VS3 shares characteristics with OPC-like cells, and its enrichment for cellular respiration and microtubule processes are in line with findings from Liu et al. who demonstrate that OPC-like cells reside in areas closer to vasculature, implying increased oxygen requirements for VS3 cells.^63^

Recently, Greenwald et al. put forth spatially defined malignant meta-programs identified across spatial GBM sections. In addition to the four Neftel et al. cell states, they identified a hypoxic MES program, a chromatin regulatory program, a proliferation and metabolism-related program, and an astrocyte-like MES program.^12^ Similarly, Zheng et al. identified a hypoxic MES module and reactive astrocyte module (in lieu of AC-like) in addition to MES-like, NPC-like, and OPC-like cells in their independent spatial analysis, which they used to predict patient prognosis and tumor aggressiveness using machine learning methods.^47^ The VS we identified share some similarities with these newer transcriptional profiles, though our correlation analysis shows concretely that they are distinct. Moreover, researchers have shown – and we have observed here – that cells in any given state tend to be spatially clustered, yet VS localization also significantly correlates with specific pathway enrichment profiles highlighting the functional significance of these states.^12,63^

In line with VS-specific enrichments and targeted GDS knockouts, VS1 demonstrated predicted sensitivities to 5-fluorouracil-derived DNA synthesis inhibitors while VS2 showed specific sensitivity to TMZ and VS3 to taxanes. VS2 sensitivity to TMZ was confirmed by analysis of scRNA-seq data of *in vivo* treated PDX from Chen et al. Additional predicted susceptibilities to antimetabolites in both VS1 and VS3 cells made it likely that gliocidin would have an effect on both of these states as well, though this was only observed for VS3. Gliocidin alone did not have a negative effect on VS1 enrichment, perhaps due to a more dose-dependent effect or resistance mechanisms in this VS. We additionally observed VS-specific compound sensitivities in both scRNA-seq and stRNA-seq data that support the *in vivo* synergy observed between CT-179 (OLIG2 inhibitor) and Depatux-M (EGFR-expressing cell-directed tubulin inhibitor) in Suter et al. (2025).^91^ Here, we observe CT-179 (VS1 targeting), microtubule inhibitors (docetaxel, paclitaxel; VS3 targeting), and EGFR inhibitors (afatinib, erlotinib; VS3 targeting) target distinct VS (Figure 7, Figures S12-S15). Taken together with our extensive characterization of the identified VS, these results validate the functional importance of each VS not only to patient treatment but to tumor integrity.

### Limitations of the study

When attempting to identify novel therapeutic targets or strategies, we desire highly specific targets that would minimize off-target effects and toxicities. Our decision to include common essential genes in our GDS contradicts this logic since common essential genes have notable impacts on cell viability across all lineages. However, the essential genes that were included a) still met our criteria of having significantly lower ES in GBM than other lineages and b) being more inclusive of genes at this step allows for more robust identification of VS downstream. On a related note, our decision to focus exclusively on the CRISPR screen data rather than the RNAi data published by DepMap was to minimize off-target effects and to ensure confidence in complete gene silencing across thousands of cell lines of varying phenotypes. Another limitation of the GDS is that the genes in it are not necessarily transcriptionally co-regulated. However, once the GDS is isolated, we believe the relative essentiality of the genes is not as important for our analysis as those genes, as a collective, being variably expressed. Nonetheless, expression of individual GDS genes is biologically important as measured by the DepMap screen (and reinforced by our work published in Rowland et al. (2025), and our approach toward VS identification using JS allowed for those co-regulated GDS genes to have a larger impact on scores. The approach toward correlation and clustering of single-cell transcriptomic data must be taken with care, and we examined alternatives to Jaccard similarity including spearman correlations, cosine similarity, and usage of a weighted scoring method. In each instance, the JS-derived clusters provided the greatest robustness, stability, and confidence in cluster validity. Moreover, given the replicability of our results across multiple scRNA- and stRNA-seq datasets, our approach and methodology demonstrate that the GDS does inform biological significance through VS.

Importantly, our VS had high concordance across all three scRNA-seq datasets used in this study. Though only 250 DEG (2.21% of all VS3 DEG) overlap between significant DEG from the Neftel, Johnson and in-house datasets, this is likely due to the relatively high number of VS3 DEG identified in the Johnson and in-house datasets (8,367 and 6,071, respectively) compared to only 634 in the Neftel dataset.

Though deconvolution methods and technologies continue to improve, none are perfect, and spatial data provides unique challenges when data are based on spots rather than single cells. Our SCDC-based deconvolution of stRNA-seq data produced relatively low proportions of VS1 detected across 18 GBM sections, leading us to use lower proportional cutoffs when analyzing correlations between spot-level VS and pathway enrichment. Despite this, our spatial enrichment and correlation results were highly congruent with our scRNA-seq derived enrichments.

### Conclusions

Transcriptional cell states are informative for modeling and predicting treatment response and resistance on the single-cell level.^14,19,30,62,68–74^ Devising effective therapies requires identification of those states and transcriptional modules specifically, over cell types, are either the most vulnerable to drug treatment or the most essential to tumor integrity. This comprehensive characterization of GBM vulnerabilities across both scRNA- and stRNA-seq data highlights the importance of multimodal approaches to drug discovery. We demonstrate that the application of essential gene signatures to expression profiling can uncover shared transcriptional regulatory modules across tumors that have actionable vulnerabilities based on extensive enrichment and correlation analyses. We thereby present a novel framework for targeted therapy development, small molecule discovery and design, and drug repurposing. VS differ in sensitivity to existing chemotherapies, and individual patient tumors can be stratified by their relative VS transcriptional profiles, allowing for the potential for tumor-specific therapeutic targeting. Gene essentiality thereby provides a framework from which we can understand and exploit tumor transcriptional profiles on the single-cell level, yielding more options for pharmacological development in the treatment of GBM.

Critically, our pipeline as presented here need not be specific to GBM, and its applicability to other cancer types holds great promise. Such applications would not only validate the robustness and versatility of our methodology, but enhance the clinical relevance and translational applicability by uncovering novel therapeutic opportunities across different malignancies. By demonstrating the efficacy of our approach and the importance of essential genes on a cell-state level across multiple cancer types, we can enhance personalized medicine and improve patient outcomes at a broad scale for other difficult-to-treat cancers.

## METHODS

### EXPERIMENTAL MODEL AND STUDY PARTICIPANT DETAILS

#### Collection of newly diagnosed and recurrent GBM tumors

Six (6) patients with GBM gave permission for tissue collection prior to tumor resection under approved IRB study 20170887 (MODCR00002196) at the University of Miami. Of the 6 enrolled patients, 3 were newly diagnosed with GBM and 3 hadd tumor recurrence after previously receiving standard of care treatment with temozolomide chemotherapy and radiotherapy. The 3 newly diagnosed patients had not received any treatment prior to resection. After surgical resection, pathologists at the University of Miami Hospital separated resected tissue into three spatially distinct groups, namely the necrotic core, enhancing edge, and infiltrating tumor. The tissue was then transferred directly from the operating suite to the laboratory on ice. Tissue from the enhancing edge was used for dissociation and scRNA-seq (see methods below).

### METHOD DETAILS

#### Identification and characterization of glioblastoma dependency signature

CRISPR loss of function screen gene effect scores in addition to cell line metadata were downloaded directly from the DepMap portal. Downloaded data were from the 23Q2 release.^20^ The post-Chronos effect scores dataset contained effect scores for 17,453 genes knocked out in 1,078 cell lines.^23^ Cell line metadata were subset to include only those lines profiled by CRISPR screen. A generalized linear regression model implemented through *limma* was then used to compare effect scores of all genes in the 49 GBM cell lines to the effect scores in all other cell line lineages (n = 1,029).^75^ Genes with significantly lower effect scores in GBM cell lines (negative effect size by linear regression and Bonferroni-adjusted p-value < 0.05, n = 377 genes) and with a mean effect score < -0.5 were selected as GBM dependency signature (GDS) genes (n = 168 genes). All analyses performed in this study were done using R and RStudio unless otherwise specified.^76^

We used the STRING database for protein-protein interaction networks and functional enrichment analysis of the GBM-specific essential genes identified from the DepMap CRISPR knockout data.^41^ All 168 genes were searched in Homo sapiens through the database’s “Multiple proteins” input. Subsequently, k-means clustering was performed using the STRING database online tool with a k of 4 to identify clusters of protein-protein interaction networks among the GBM essential genes. Additional characterization of GDS genes via over-representation analysis was performed as described in **Quantification and Statistical Analysis**.

#### External single-cell RNA sequencing data set acquisition and processing

Previously published data from Neftel et al. was downloaded from GEO (GSE131928) as processed, log-normalized counts.^8^ These data were then subject to our in-house pipeline for scRNA-seq processing using R and *Seurat*.^29,77^ The log-normalized counts matrix containing expression data for 26 tumors was converted to a *Seurat* object and downloaded metadata from the authors added. Six (6) pediatric GBM tumors were excluded from downstream analysis at this point, leaving a total of 20 GBM tumors. Individual cells were confirmed to have < 20% mitochondrial transcripts and contain data on more than 100 features. Subsequently, cells were split by individual tumor and subject to normalization and variance stabilization using a regularized negative binomial regression implemented within the *scTransform* R package.^29,78^ Individual tumor scRNA-seq data were subsequently integrated using an anchor-based technique utilizing the Pearson residuals.^29,77^ Normalized, integrated values were then used for shared-nearest-neighbor (SNN) clustering, dimensional reduction, cell type verification, and cell type identification. Cells were scored for cell cycle phase and mitochondrial RNA content using *Seurat* and expression data were scaled with these variables regressed out. Downstream analysis including differential expression and enrichment scoring was performed using log-normalized counts (see below).

Previously published data from Johnson et al. was downloaded from Synapse (https://synapse.org/singlecellglioma) and processed using our in-house scRNA-seq pipeline as described above.^30^

Data from Jackson et al. (2025) were downloaded from GEO (GSE278456) as a pre-processed *Seurat* .RDS file.^31^ Data were evaluated for quality control metrics as described above. For analysis of GDS expression between central nervous system (CNS) neoplasms, single-cells were module-scored for GDS genes, which were then normalized to normal brain GDS MS for each tumor. The mean MS was calculated for each tumor and comparisons were performed using a one-way ANOVA with Tukey’s post-hoc test. Statistical analysis and comparisons for astrocytoma were not included as only 1 astrocytoma tumor was included in the data set (Figure S1F).

Data from Chen et al. (2024) were downloaded from GEO (GSE277301) as a preprocessed *Seurat* .RDS file.^59^ Data were evaluated for quality control metrics as described above. This dataset contains data for 12 GBM PDX orthotopically implanted in mice and treated with TMZ, gliocidin, a combination of TMZ and gliocidin, or DMSO (3 mice per treatment group). For comparisons between treatments, single-cell enrichments for vulnerability state (VS) expression were calculated using *singscore* and normalized to DMSO-treated tumors. Importantly, tumors from the same treatment group were pooled prior to scRNA-seq. VS enrichment scores for each treatment group were normalized to DMSO treatment and the number of cells per treatment group. Single-cell level differences between VS normalized enrichments were calculated using one-way ANOVA with pairwise significance determined using Tukey’s post-hoc test and significance set at p < 0.05.

#### Single-cell RNA sequencing of patient glioblastoma tumors

The 6 patient GBM samples described in **Experimental Model and Study Participant Details** were dissociated using the Worthington Biochemical Papain Dissociation System.^79^ Tumor tissue was finely minced with a sterile surgical blade and incubated in sterile Earl’s Balanced Salt Solution (EBSS) containing 20 units of papain per mL in 1mM L-cysteine with 0.5 mM EDTA and DNase for 45 minutes at 37°C while shaking. Following papain incubation, the tissue was gently triturated with a 5 mL serological pipette until it was adequately dissociated. The resulting suspensions were pelleted and resuspended in EBSS with ovomucoid protease inhibitor, bovine serum albumin (BSA), and DNase. This suspension was layered on top of ovomucoid protease inhibitor solution with BSA to form a gradient, which was centrifuged at 70xg for 6 minutes. The supernatant containing cellular debris was then removed. The pellets were subsequently resuspended, and the gradient centrifugation was repeated. The resulting pellet was then washed 2 times with ice-cold PBS with 0.1% BSA, passed through 0.45 uM cell strainer, and resuspended in ice-cold PBS with 0.1% BSA. This final suspension was then checked for cell viability by staining an aliquot with acridine orange and propidium iodide (AO/PI) and counted on a Nexcelom K2 Cellometer. All samples had cell viability greater than 90% prior to loading on the single-cell chips.

Single cells were captured using a 10X Genomics Chromium Controller, and patient single-cell gene expression libraries were prepared using 10X Genomics 5’ Gene Expression chemistry. All libraries were sequenced on an Illumina NextSeq500. Alignment and demultiplexing was performed using CellRanger, and filtered UMI matrices were used for downstream analysis using the *Seurat* package.^29^ Within *Seurat*, poor quality cells were further filtered by removing outliers based on the number of detected UMIs and unique transcripts, as well as percent expression of mitochondrial transcripts. Pass-filter cells were those with greater than 200 and less than 4,500 detected features, and with less than 12.5% mitochondrial transcript expression. Filtered data were normalized and variance stabilized using a regularized negative binomial regression implemented within the *scTransform* R package, and individual tumor scRNA-seq datasets were subsequently integrated with an anchor-based technique utilizing the Pearson residuals.^29,77^ Normalized, integrated values were then used for initial shared-nearest-neighbor (SNN) clustering, dimensional reduction, and cell type identification. Downstream analysis including differential expression and enrichment scoring was performed using log-normalized counts.

#### Discovery of glioblastoma vulnerability states

The processed Neftel scRNA-seq dataset was split by patient tumor (n = 20 adult GBM) and the scaled, log-normalized expression of the 168-gene GDS genes was binarized. For each independent tumor, scaled expression of a gene > 0 was assigned “1” and all other values were assigned “0”. The Jaccard Similarity Index (Jaccard Index, Jaccard Similarity, JS) was then computed between pairs of single cells based on these binarized expression values to create a pairwise cell-cell JS matrix for each tumor. The agglomerative coefficient was then computed for clustering methods including ward, average, single, and complete clustering. For each tumor, the ward clustering method had the highest agglomerative coefficient and was chosen for all subsequent clustering.

The JS matrix for each tumor was then clustered by Ward D2 hierarchical clustering for all k between 2 and 30. For each value of k, clusters were excluded if it included fewer than 5 cells or greater than 80% of the total number of neoplastic cells in the tumor to ensure high quality differential expression analysis (downstream). After exclusions, differential expression was performed between all clusters for each k using *Seurat’s FindMarkers* with a log2FC threshold set to > 0 using MAST statistical testing.^42^ Clusters at each k were retained if they contained greater than 50 genes (N50) with an adjusted p-value < 0.05 and at least 1 gene (N1) with an adjusted p-value < 0.001.

After clustering on each independent tumor, JS were calculated between retained clusters using the defined N50 genes (modules) for each cluster. For pairs of modules with > 75% similarity (JS > 0.75), we retained the larger module (cluster with the highest number of genes in N50). This resulted in 25 modules identified across 11 tumors, which were subsequently subjected to JS and Ward D2 hierarchical clustering with k = 3, uncovering 3 vulnerability states (VS). Cluster stability and robustness were confirmed by Calinski-Harabasz index and *clustree* visualizations (Figure S5).^80^ All heatmaps were visualized using the *pheatmap* R package.^81^

#### Single-cell vulnerability state signature identification and cell state assignment

Signature genes for each identified VS were identified by aggregating the top 100 genes across all modules comprising the VS, ranked by log2FC. Genes with more than one instance within a single signature were aggregated by taking the mean log2FC prior to this, resulting in 100-gene signatures per VS, consisting of 292 unique genes across all VS.

VS signature genes were used for VS assignment within the Neftel, Johnson, and Suter datasets. This was done using *singscore*, an independent-sample gene-set scoring method implemented in R, to score each neoplastic single-cell for ranked expression of the 300 marker genes (Figure S4), binned according to VS (100 genes per VS).^43,44^ Neoplastic cells were then assigned to a VS based on which VS marker gene-set had the highest *singscore* enrichment.

Any additional gene set scoring was performed using the Module Score (MS) calculation provided by *Seurat*.^29^ MSs provide a scaled method of evaluating single-cell level expression of a collection of genes.^29,82^

#### Spatial transcriptomic data acquisition, processing, deconvolution, and analysis

10X Visium spatial transcriptomic RNA sequencing (stRNA-seq) data was acquired from Ravi et al. (2022) as h5 counts with corresponding H&E-stained section images from Dryad.^15,83^ These data were processed and analyzed using *Seurat* as described above for scRNA-seq data. Of 28 total sections, 18 are GBM tumor sections; 8 of 28 are surgical entry site normal cortex, and the 2 remaining samples are IDH-mutant and thereby not classified a GBM and not analyzed here. In short, poor quality spots without counts data were removed, and filtered data were normalized and variance stabilized image-by-image using scTransform.^29,78^ Individual images were subsequently merged and integrated using a canonical correlation analysis (CCA)-based method implemented through *Seurat*. Normalized, integrated values for all spots across all sections were then used for SNN clustering and dimensionality reduction. Downstream analyses including differential expression and enrichment were performed on scaled data regressed for cell cycle genes and mitochondrial RNA.

For spot-based deconvolution of stRNA-seq data, we used *SCDC*, an RNA-seq deconvolution method that can leverage cell-type specific gene expression information from multiple scRNA-seq references to address batch-effect confounding.^48^ Raw counts of the Ravi et al. stRNA-seq data were used for spot-based deconvolution with our in-house scRNA-seq dataset used as the reference set to provide single-cell type annotations. Visualization of deconvolution results from *SCDC* was performed with *SPOTlight* to represent each spot as a pie chart with each cell type predicted at that spot colored as a pie section (Figure S9).^48,84^

#### Single-cell regulatory network inference and clustering (SCENIC) analysis

*SCENIC* analysis using our in-house dataset was performed using the pySCENIC implementation and python through a Docker image.^50^ The raw expression matrix of all single-cells were used as input to calculate predicted transcription factor regulon activities. Resulting regulon activity scores were then aggregated between each VS and compared using a GLM implemented through *limma* in R.^75^ Significantly different regulon activities were defined as those with p < 0.05 between VS.

#### LINCS L1000 CRISPR knockout transcriptional consensus signature analysis

The Library of Integrated Network-Based Cellular Signatures (LINCS) L1000 dataset was screened for CRISPR knockouts to identify gene knockouts that overlapped with the GDS. The L1000 assay contains 978 “landmark” genes that can be measured to algorithmically infer the expression of 11,300 additional genes.^51,52^ Transcriptomic data from cell lines perturbed by those knockouts was then analyzed using a previously-published algorithm to identify transcriptional consensus signatures (TCS) of L1000 compounds, representing the top upregulated and downregulated genes in response to perturbation.^51–54^ This method was adopted for CRISPR knockouts as the perturbation of interest instead of small molecules. Transcriptional consensus signatures for each CRISPR knockout (39 of 168 GDS genes) were identified using this algorithm and subsequently scored for enrichment in our in-house, Johnson data, and Neftel scRNA-seq datasets independently. Using *singscore*, each TCS was scored for discordance with the ranked transcriptomic profile of each single-cell with each TCS split by up- and down-regulated genes. CRISPR perturbation signature (XPS) scores for each neoplastic single-cell were then aggregated first by VS and across datasets to calculate a combined XPS score.

XPS scores were then inverted such that predicted sensitivity to CRISPR knockout was represented by a positive value.

#### Inferred cell sensitivity operating on the integration of single-cell expression and L1000 expression signatures (ISOSCELES) and drug connectivity analysis

Analysis of predicted single-cell level small molecule sensitivities was conducted using ISOSCELES (Suter et al. 2025, *bioRxiv*).^91^ This tool uses the principle of disease signature reversal to perform these calculations. Our in-house scRNA-seq dataset underwent ISOSCELES analysis through the Shiny implementation of the application with untransformed cells identified as non-neoplastic cells of the TME and the comparison groups as neoplastic cells split by VS. The resulting consensus correlations to L1000 TCS for each single cell were then downloaded and analyzed using R. Unless otherwise specified, consensus correlations (drug connectivities) were inverted so that higher scores (more positive) represented predicted compound sensitivities. Subsequently, single-cell level correlations were compared between VS using a GLM implemented through *limma* in R.^75^ Significantly different compound sensitivities were defined as those with BH-adjusted p < 0.05 between VS. L1000 compounds were subset to only include analysis of drugs and small molecules annotated as FDA approved for treatment of oncologic diseases.

### QUANTIFICATION AND STATISTICAL ANALYSIS

#### Module Score and single-cell enrichment comparisons and statistical analysis

All comparisons between single-cells or tumors that used MS or single-cell enrichment scores were performed by one-way ANOVA using a Tukey’s post hoc test for individual pairwise differences. Resulting p-values were corrected using a Bonferroni adjustment with significance set to p < 0.05.

#### Differential expression and differential enrichment analysis

All differential expression analyses were performed using *Seurat’s FindMarkers* function with a baseline log2FC cutoff > 0 and statistical testing performed with MAST.^29,42^ Significance was determined by MAST-adjusted p-values < 0.05.

Differential enrichment or over-representation analyses were performed using the Molecular Signatures Database (MSigDB) with the specific collections named in each instance. Notable collection subsets include KEGG, KEGG MEDICUS, WikiPathways, and Reactome. R packages used for these analyses include *enrichR*, *msigdbr*, and *ClusterProfiler*.^32,85,86^ In each instance, significant differences were those with Benjamini Hochberg (BH)-adjusted p-values < 0.05 by Fisher’s exact test.

Visual comparisons between DEG profiles between datasets were performed using the *VennDiagram* R package to highlight overlapping DEG for each VS.^87^

#### Cell and tumor type classification correlation analysis

Transcriptional signatures from previously-published GBM classification systems were acquired from Verhaak et al. (2010), Neftel et al. (2019), and Ravi et al. (2022).^8,15,45^ Each single-cell in our in-house scRNA-seq dataset was scored for enrichment of these signatures, in addition to our VS signatures, using *singscore* as mentioned in **Method Details**. We then calculated Spearman correlations on the resulting single-cell enrichment scores to calculate and subsequently visualize with *corrplot* similarities between transcriptional classifiers.^88^

#### Gene set co-regulation analysis and spot-level correlation analysis

The 18 GBM tumor spatial sections were subject to gene set co-regulation analysis (GESECA) using the *fgsea* R package and the Gene Ontology Biological Processes (GOBP).^49,89,90^ Input data were the results of reverse principal component analysis integration performed on scaled RNA-seq data from the Spatial assay with cell cycle genes and mitochondrial RNA regressed out. Minimum gene set size was set to 2, maximum size was set to 500. Enriched pathways were filtered for those with BH-adjusted p-values less than 0.05 by Fisher’s exact test, and then the top 50 by adjusted p-value were visualized using the z-score of the GESECA enrichment score.

In addition to spot-level proportion data on VS from *SCDC* deconvolution, we used *singscore* to calculate an enrichment score for each VS at each spot across all 18 tumor sections. Similarly, we used *singscore* to calculate enrichment scores for significantly enriched GOBP terms to allow for accurate correlation comparisons between VS and GOBP pathways. We then correlated VS and GOBP enrichment scores for all spots across all 18 sections. For each VS, we correlated those spots with > 0% (VS1) or > 50% (VS2 and VS3) content in any given spot with GOBP enriched terms using *singscore*-based enrichment scores. Different proportional cutoff values were used for each VS given the relatively low levels of VS1 detected across all GBM sections in the stRNA-seq data to include similar numbers of spots in each independent correlation analysis. Correlations greater than 0.5 (VS1 and VS2) or 0.3 (VS3) are available in File S3.

#### SCENIC and ISOSCELES single-cell differential score analysis

SCENIC analysis as described in **Method Details** provides regulon activity scores for each single-cell in our in-house scRNA-seq dataset. Single-cell-level differences in regulon activity scores was calculated between VS using a GLM implemented through *limma*.^75^ Regulons with pairwise, Bayes-corrected and Bonferroni-adjusted p-values < 0.05 between VS were used for subsequent analysis and visualization (Figure 6B). Pseudobulk regulon activity scores for all cells belonging to a single VS were calculated using the mean for all cells belonging to a VS. Scaling across pseudobulk scores for each regulon was performed for visualization.

Similarly, ISOSCELES analysis as described in **Method Details** provides drug connectivity scores for each single-cell in our in-house scRNA-seq dataset. Single-cell-level differences in drug connectivity between VS were calculated using a GLM implemented through *limma*.^75^ Drugs with pairwise, Bayes-corrected and Bonferroni-adjusted p-values < 0.05 between VS were used for subsequent analysis and visualization (Figure 7A). Pseudobulk connectivity scores for all cells belonging to a single VS were calculated using the mean for all VS cells. Scaling across pseudobulk scores for each drug was performed for visualization.

#### LINCS L1000 CRISPR knockout perturbation response analysis

The Library of Integrated Network-Based Cellular Signatures (LINCS) L1000 dataset was screened for to identify CRISPR knockouts that intersected with GDS genes. The L1000 assay contains 978 “landmark” genes that can be measured to algorithmically infer the expression of 11,300 additional genes.^51,52^ Transcriptomic data from cell lines perturbed by those knockouts was then analyzed using a previously-published algorithm to identify transcriptional consensus signatures (TCS) of L1000 compounds, representing the top upregulated and downregulated genes in response to perturbation.^51–54^ This method was adopted for CRISPR knockouts as the perturbation of interest instead of small molecules.

Transcriptional consensus signatures for each CRISPR knockout (39 of 168 GDS genes) were identified using this algorithm and subsequently scored for enrichment in our in-house, Johnson data, and Neftel scRNA-seq datasets independently. Using *singscore*, each TCS was scored for discordance with the ranked transcriptomic profile of each single-cell with each TCS split by up-and down-regulated genes. CRISPR perturbation signature (XPS) scores for each neoplastic single-cell were then aggregated by mean, first by VS and second across datasets to calculate a combined XPS score. XPS scores were then inverted (multiplied by -1) such that predicted sensitivity to CRISPR knockout was represented by a positive value.

#### Patient derived xenograft in vivo treatment comparisons and statistical analysis

The Chen et al. (2024) dataset contains scRNA-seq data for 12 GBM PDX orthotopically implanted in mice and treated with TMZ (200μM), gliocidin (5μM), or a combination of TMZ and gliocidin, or DMSO (3 mice per treatment group) for 10 days. scRNA-seq of tumors collected at mouse endpoints are described in their publication.^59^ Importantly, tumors from the same treatment group were pooled prior to scRNA-seq. For comparisons between treatments, single-cell enrichments for vulnerability state (VS) expression were calculated using *singscore*. VS enrichment scores for each treatment group were normalized to DMSO treatment and the number of cells analyzed per treatment group. Single-cell level differences between VS normalized enrichments were calculated using one-way ANOVA with pairwise significance determined using Tukey’s post-hoc test and significance set at p < 0.05.

## ADDITIONAL RESOURCES

ISOSCELES Shiny Web GUI: https://robert-k-suter.shinyapps.io/isosceles^91^

ISOSCELES R package: https://github.com/AyadLab/ISOSCELES^91^

## RESOURCE AVAILABILITY

### Data and code availability

All data and code needed to generate and reproduce the materials and conclusions presented in this manuscript are available in the body of the text, the Supplemental Materials, and/or the Ayad Laboratory GitHub repository (https://github.com/AyadLab/scZeroDay). Data, code, and materials are made available to any researcher. Our generated single-cell data of patient GBM tumors, including processed Seurat objects, are available via the Gene Expression Omnibus (GEO) with accession ID GSE229779 (Reviewer Token: yzghmqawxbodtmf). Neftel et al. (2019) scRNA-seq data were obtained from GEO with ID GSE131928. Johnson et al. (2022) scRNA-seq data were obtained from Synapse at https://synapse.org/singlecellglioma. Spatial transcriptomic data published by Ravi et al. were obtained from Dryad at https://doi.org/10.5061/dryad.h70rxwdmj. Data from Jackson et al. (2025) were downloaded from GEO with ID GSE278456. Data from Chen et al. (2024) were downloaded from GEO with ID GSE277301. Any additional information required to reanalyze the data reported in this paper is available from the lead contact upon request.

## ACKNOWLEDGMENTS

We would like to thank Dr. Siddharth Jain for his helpful comments and advice. This work has been supported by BellRinger at the Georgetown University Lombardi Comprehensive Cancer Center (LCCC). We would like to thank LCCC and the Georgetown University Tumor Biology Program. We additionally thank all other members of the Ayad laboratory for invaluable discussion of this research.

## AUTHOR CONTRIBUTIONS

**Conceptualization**: N.G.A., R.K.S., M.S., M.D. **Methodology, investigation, visualizations**: M.D. **Single-cell RNA sequencing samples and data**: R.K.S, M.E.I, R.J.K. **Computational design**: M.D., R.K.S., M.S., R.C. **Data analysis**: M.D. **Writing, editing, and revisions**: M.D., R.K.S., N.G.A. **Supervision**: N.G.A., R.K.S.

## DECLARATION OF INTEREST

The authors declare no financial, personal, or ethical conflicts of interest. This content is solely the responsibility of the authors and does not necessarily represent the official views of the National Institutes of Health.

## FUNDING INFORMATION

Research reported in this publication was supported by the National Institute of Neurological Disorders and Stroke (NINDS) of the National Institutes of Health under grand number RM1NS133003 and by Georgetown University Lombardi Comprehensive Cancer Center (LCCC) BellRinger.

## Notes

### Competing Interest Statement

The authors have declared no competing interest.

### Summary of Updates

Updated funding information in submission portal and manuscript

