## Supplementary material for "Identification of disease-specific vulnerability states at the single-cell level": Figures S1-S11, Table S1

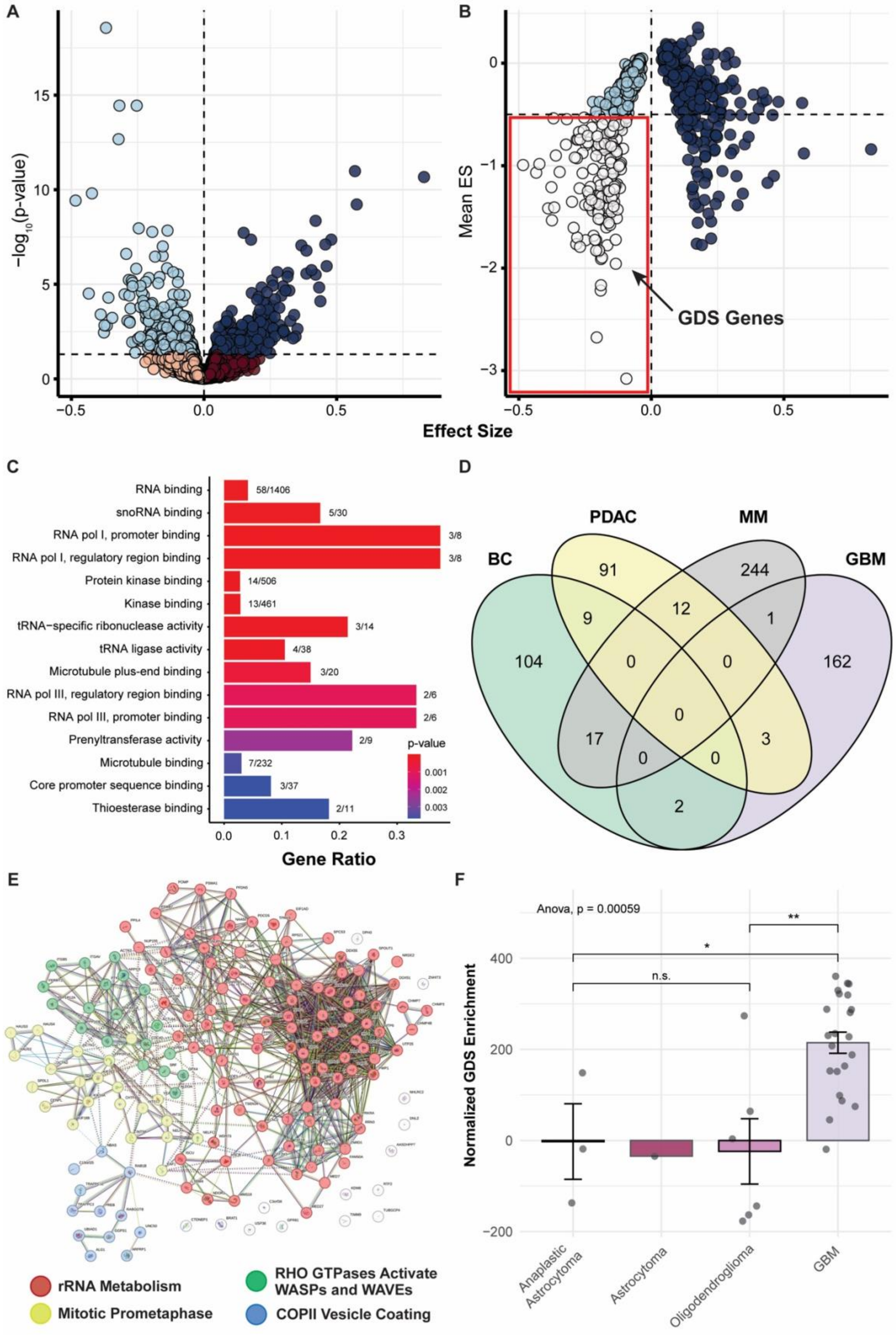

A
B
−log10(p-value)
Mean ES
GDS Genes
Effect Size
C
RNA binding 58/1406
snoRNA binding 5/30
RNA pol I, promoter binding 3/8
RNA pol I, regulatory region binding 3/8
Protein kinase binding 14/506
Kinase binding 13/461
tRNA−specific ribonuclease activity 3/14
tRNA ligase activity 4/38
Microtubule plus-end binding 3/20
RNA pol III, regulatory region binding 2/6
RNA pol III, promoter binding 2/6
Prenyltransferase activity 2/9
Microtubule binding 7/232
Core promoter sequence binding 3/37
Thioesterase binding 2/11
p-value
Gene Ratio
D
BC
PDAC
MM
GBM
E
rRNA Metabolism
Mitotic Prometaphase
RHO GTPases Activate WASPs and WAVEs
COPII Vesicle Coating
F
Anova, p = 0.00059
n.s.
Normalized GDS Enrichment
Anaplastic Astrocytoma
Astrocytoma
Oligodendroglioma
GBM

**Figure S1. Further characterization of glioblastoma dependency signature**

(**A**) Scatter plot depicting the differences in gene ES in GBM-lineage cell lines (n = 49) compared to all other cell lines (n = 1,029) by a linear model using all 17,453 genes profiled by DepMap's CRISPR screens. Negative effect size indicates genes with lower effect scores in GBM cell lines. The negative log of the adjusted p-value is represented on the y-axis, with a p-value cutoff of 0.05 represented by the horizontal dashed line. Blue coloring (both shades) represents genes with a significant p-value (< 0.05). Candidate GBM-specific essential genes are those with p < 0.05 and negative effect size (light blue). (**B**) Scatter plot depicting the second level of selection for the GBM dependency signature (GDS). All genes with adjusted p-value < 0.05 from (A) were plotted as their mean ES in all GBM-lineage cell lines (y-axis) versus their effect size. Identified GDS genes are those with a negative effect size and a mean effect score < -0.5 (white). (**C**) Additional enrichment analysis of the 168 genes that comprise the GDS, conducted using MSigDB's Gene Ontology Molecular Function (GOMF) gene sets. Color indicates BH-adjusted p-value. Gene ratio (x-axis) is the ratio of genes present in the GDS to the genes present in the named GOMF gene set (fractional representation to the right of each bar). (**D**) Four-way Venn diagram depicting the number of overlapping dependency genes identified for multiple cancer cell line lineages using methods identical to those used for GDS identification. (BC = breast cancer, PDAC = pancreatic ductal adenocarcinoma, MM = multiple myeloma). (**E**) STRING network analysis highlighting protein-protein interactions among the 168 GDS genes. Colors represent the GO-annotated results of k-means clustering with k = 4. (**F**) GDS enrichment scores by tumor (black dots) across different CNS neoplasms. Enrichment scores were normalized to normal brain. Data derived from Jackson et al. (2025) scRNA-seq data. Statistical analysis performed by one-way ANOVA with Tukey's post-hoc test (* p < 0.05, ** p < 0.01, n.s. = not significant).

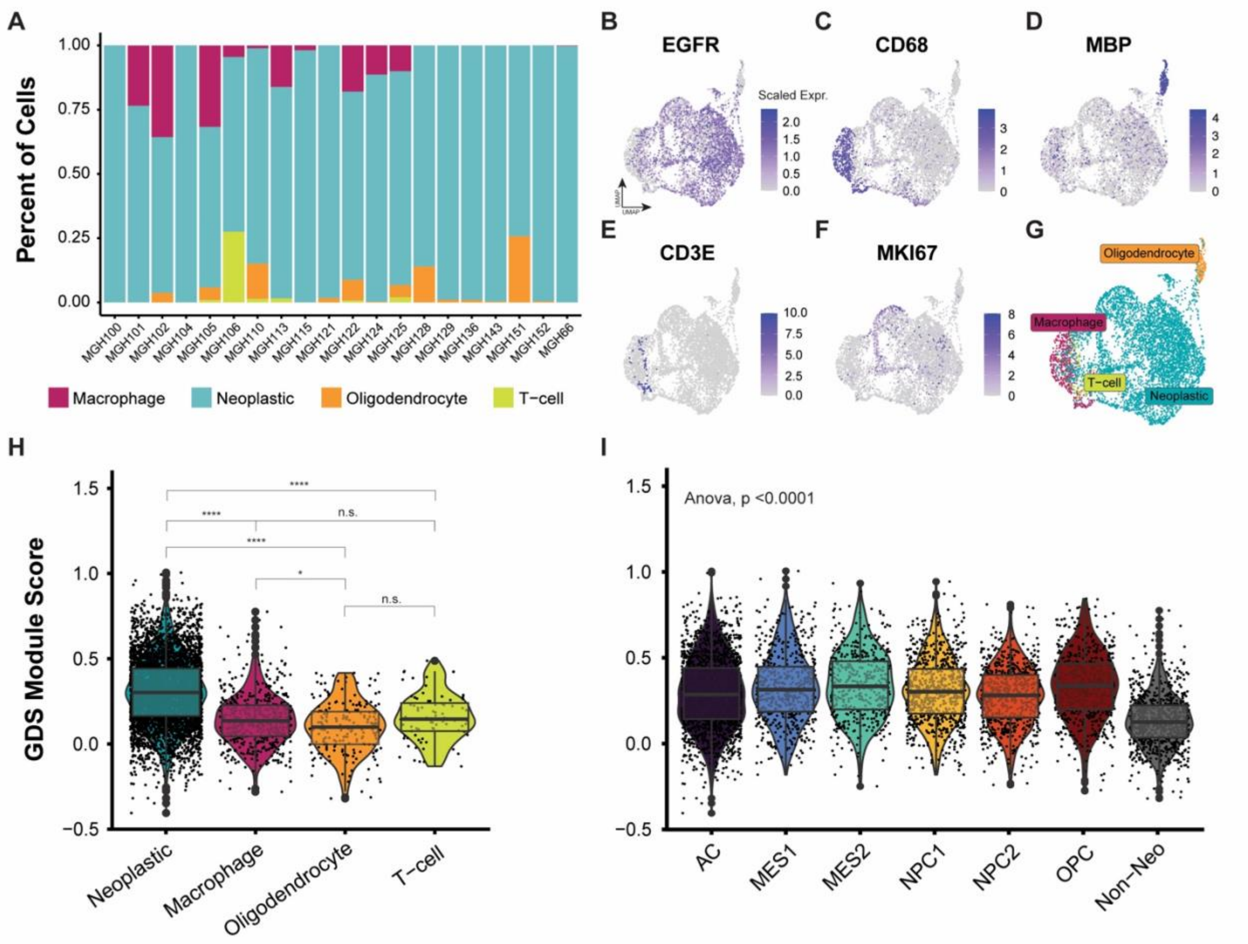


**Figure S2. Characterization of 20 adult glioblastoma tumors subset from Neftel et al. scRNA-seq dataset**

(**A**) Stacked bar plot showing the proportion of cell types in the n = 20 adult GBM tumors present in the Neftel et al. (2019) scRNA-seq dataset. (**B**) UMAP plot of the Neftel et al. (2019) scRNA-seq dataset with cell type labels. (**C-G**) UMAP plots depicting scaled expression of named cell type lineage markers. An expression value of 0 includes cells with negative scaled expression. (**H**) Boxplot with underlying violin plot that shows the MS of the GDS in each single cell (black dots) in the Neftel et al. scRNA-seq dataset, grouped by cell type. Cell type MS are compared by an ANOVA with Tukey's post-hoc test for multiple comparisons. (**I**) Boxplot with underlying violin plot that shows the MS of the GDS in each single cell (black dots) in the Neftel et al. scRNA-seq dataset, grouped by Neftel cell states. Cell state GDS MS are compared by an ANOVA with Tukey's post-hoc test (**** = $p < 0.0001$, *** = $p < 0.001$, ** = $p < 0.01$, * = $p < 0.05$, n.s. = not significant).

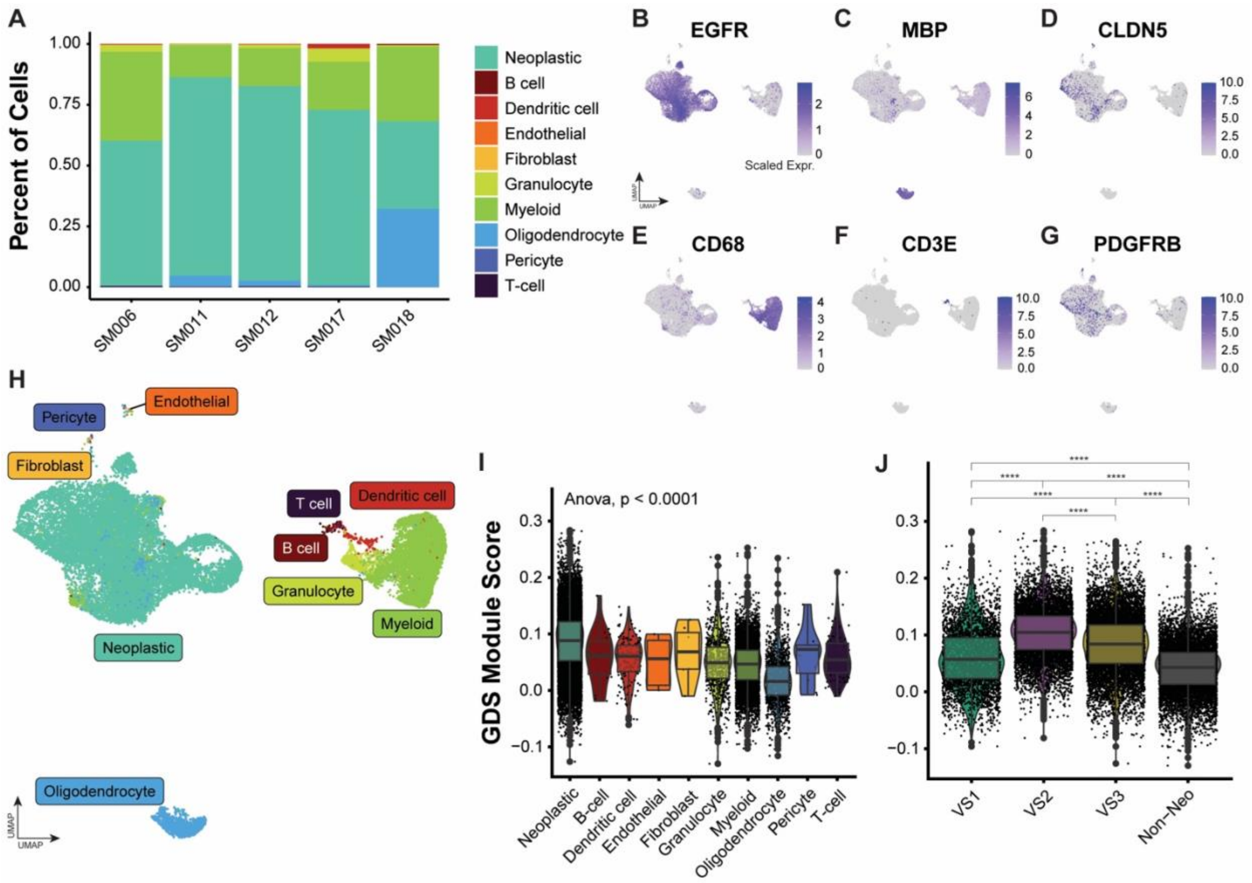


**Figure S3. Characterization of 5 adult glioblastoma tumors subset from Johnson et al. scRNA-seq dataset**

(**A**) Stacked bar plot showing the proportion of cell types present in the Johnson et al. (2021) scRNA-seq dataset. (**B-G**) UMAP plots depicting scaled expression of named cell type lineage markers. An expression value of 0 includes cells with negative scaled expression. (**H**) UMAP plot of the Johnson et al. (2021) scRNA-seq dataset with cell type labels. (**I**) Boxplot with underlying violin plot that shows the MS of the GDS in each single cell (black dots) in the Johnson et al. scRNA-seq dataset, grouped by cell type. Cell type MS are compared by an ANOVA with Tukey's post-hoc test. (**I**) Boxplot with underlying violin plot that shows the MS of the GDS in each single cell (black dots) in the Johnson et al. scRNA-seq dataset, grouped by VS assignment. Cell state MS are compared by an ANOVA with Tukey's post-hoc test (**** = p < 0.0001, *** = p < 0.001, ** = p < 0.01, * = p < 0.05, n.s. = not significant).

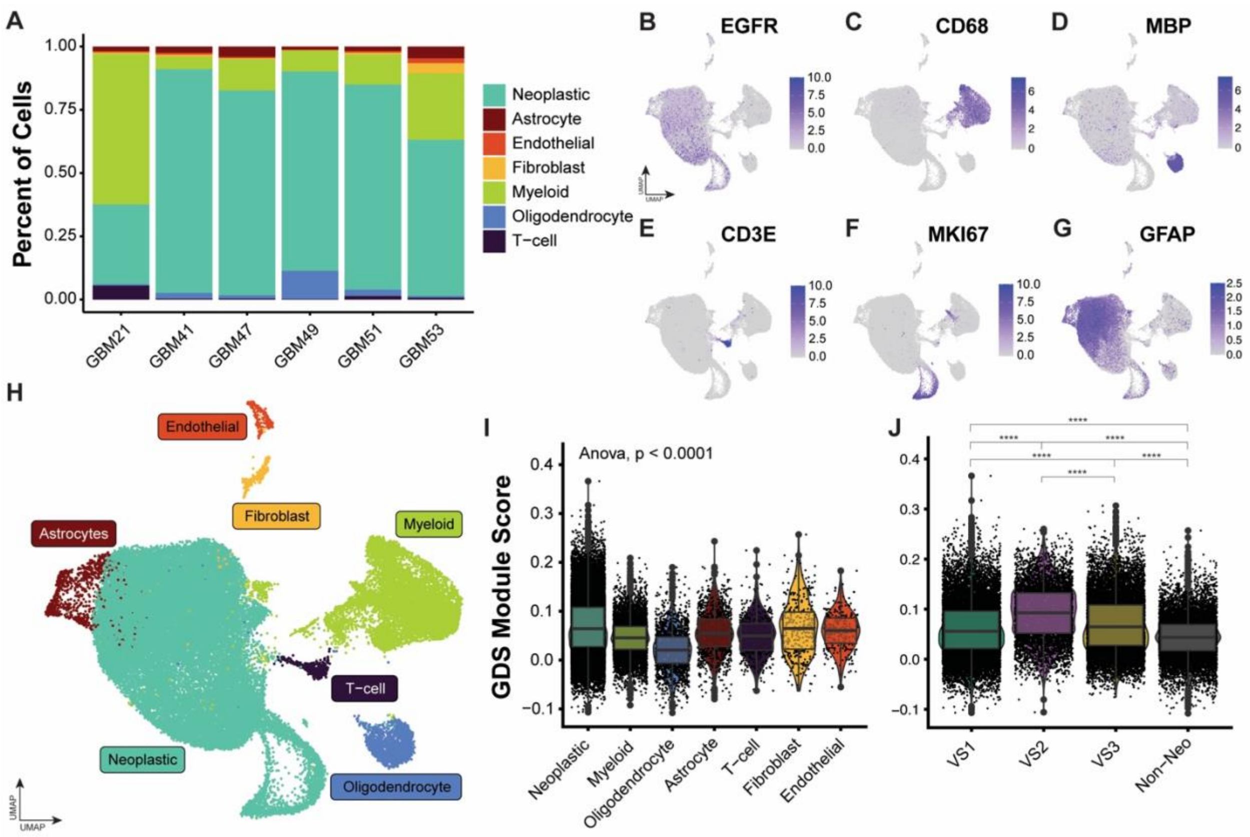


**Figure S4. Characterization of 6 adult glioblastoma tumors by scRNA-seq**

(**A**) Stacked bar plot showing the proportion of cell types identified in our in-house scRNA-seq dataset. (**B-G**) UMAP plots depicting scaled expression of named cell type lineage markers. An expression value of 0 includes cells with negative scaled expression. (**H**) UMAP plot of our in-house scRNA-seq dataset with identified cell type labels. (**I**) Boxplot with underlying violin plot that shows the MS of the GDS in each single cell (black dots) in our in-house dataset, grouped by cell type. Cell type MS are compared by an ANOVA with Tukey's post-hoc test. (**I**) Boxplot with underlying violin plot that shows the MS of the GDS in each single cell (black dots) grouped by VS assignment. Cell state MS are compared by an ANOVA with Tukey's post-hoc test (**** = p < 0.0001, *** = p < 0.001, ** = p < 0.01, * = p < 0.05, n.s. = not significant).

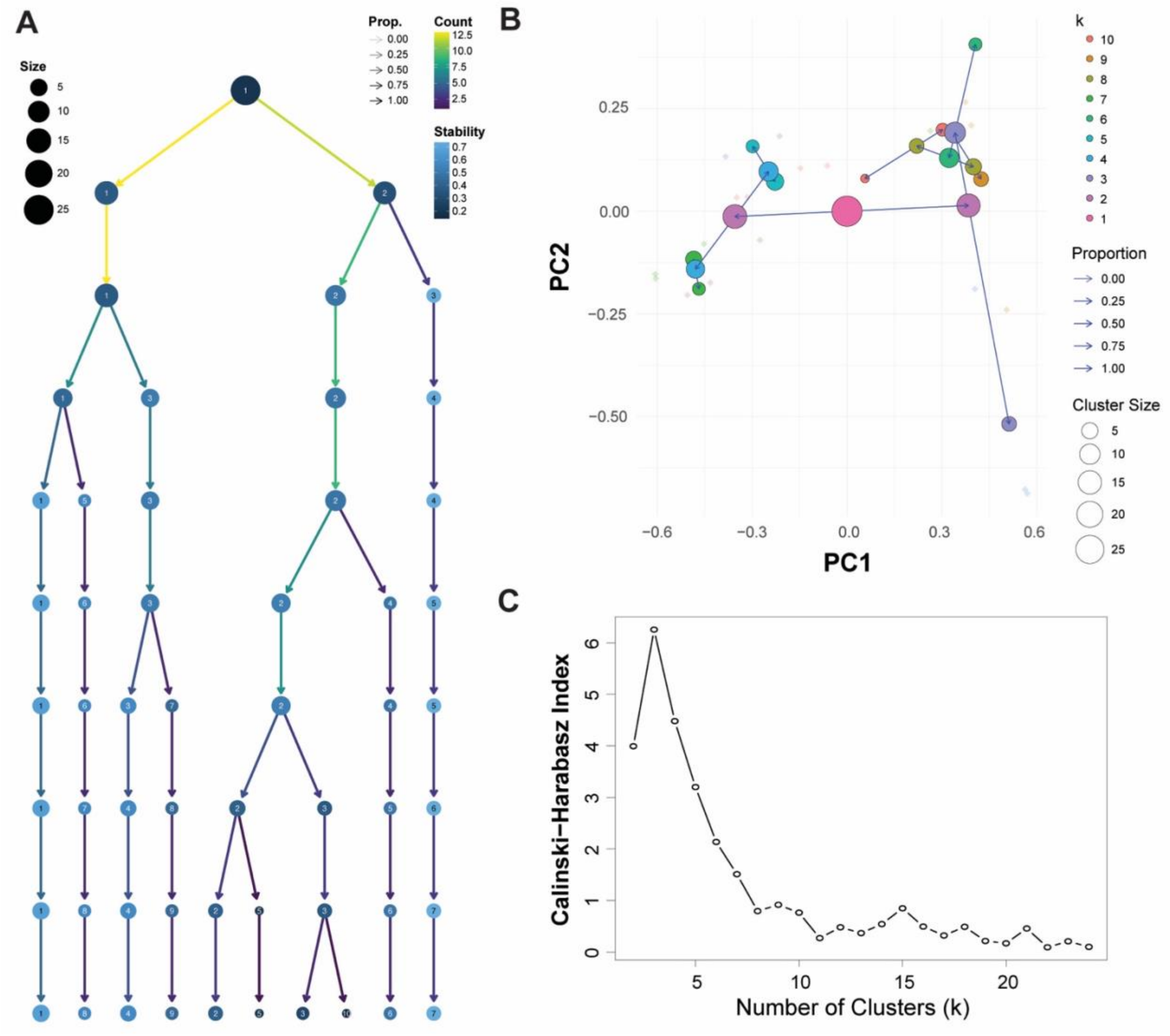


**Figure S5. Vulnerability state clustering validation**

(**A**) Clustree branching diagram displaying hierarchical clusters at different values of k. Each row represents a different k value, and each dot represents a numbered cluster at that k. Dot size indicates the number of modules assigned to that cluster. Arrow color represents the number of modules transitioning between clusters between k values. Dot color represents cluster stability. (**B**) Scatter plot displaying PC1 and PC2 of the JS matrix, overlayed with *clustree* diagram. (**C**) Calinski-Harabasz index plot for determining optimal k for hierarchical clustering.

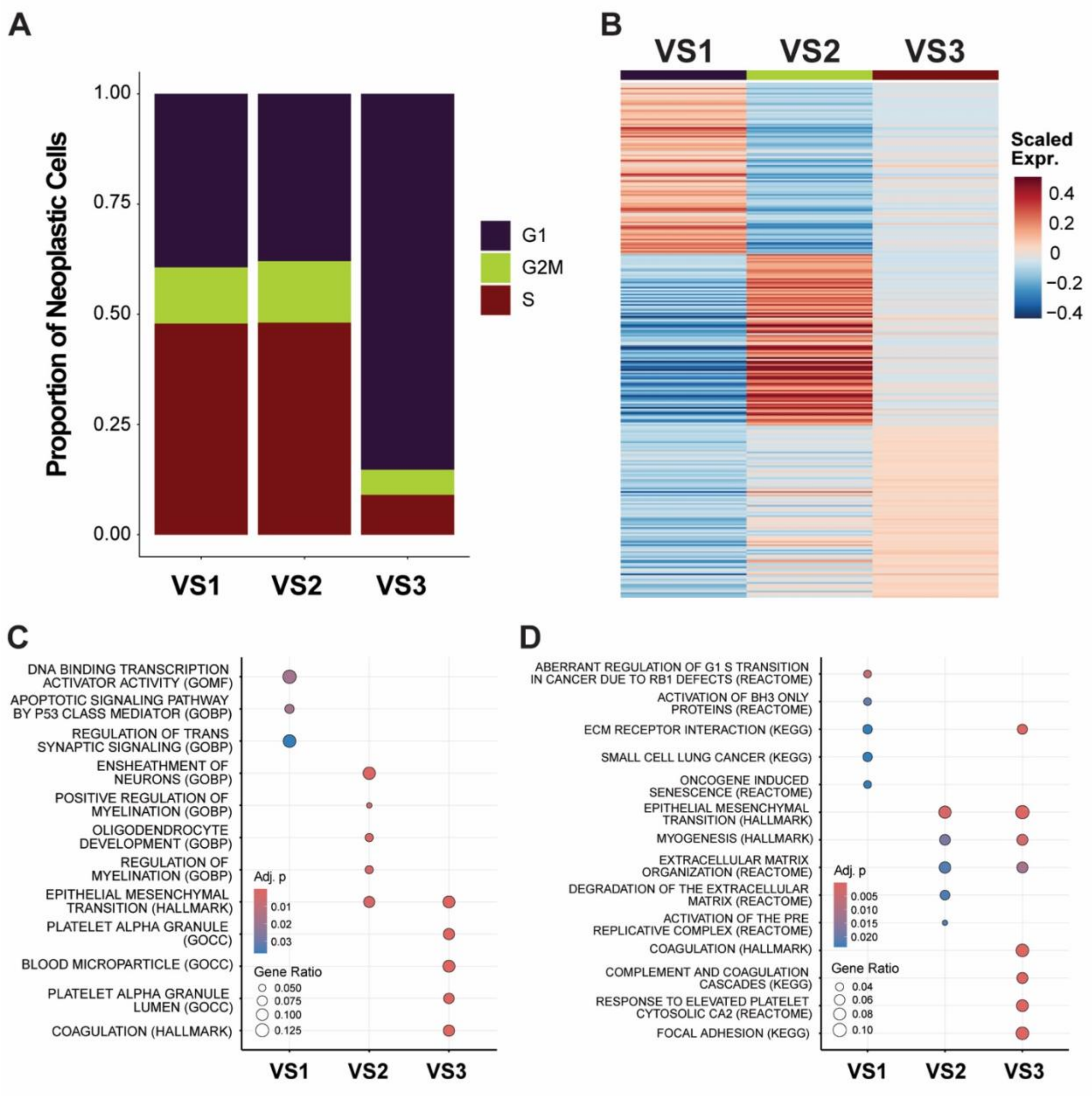


**Figure S6. Additional characterization of GBM vulnerability states**

(**A**) Proportion of neoplastic single cells at each phase of the cell cycle, grouped by VS. (**B**) Heatmap showing the pseudo-bulk scaled expression of the top 100 DEG between VS in the Neftel et al. (2019) dataset. (**C**) The top 5 differentially enriched terms between the top 100 DEG per VS using MSigDB's C5 and H collections. (**D**) The top 5 differentially enriched terms between the top 100 DEG per VS using MSigDB's C2 and H collections.

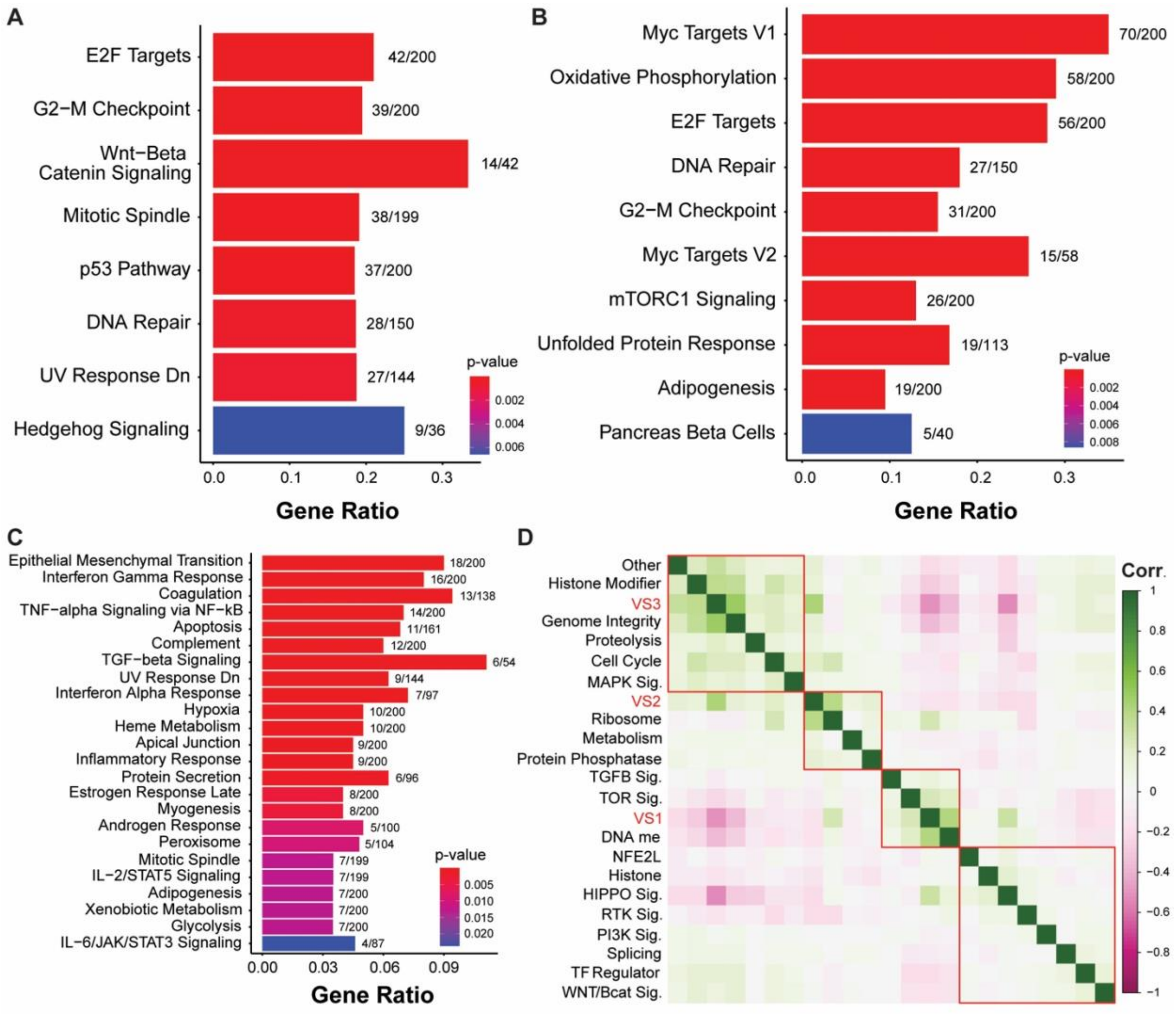


**Figure S7. Vulnerability state differentially expressed genes are conserved across datasets**

(**A-C**) Bar plots depicting the top enriched pathways for overlapping VS DEG between all three scRNA-seq datasets (Neftel, Johnson, In-House) with (**A**) corresponding to VS1 enrichments, (**B**) corresponding to VS2 enrichments, and (**C**) corresponding to VS3 enrichments. Enrichment analysis was performed using enrichR and the MSigDB Hallmarks gene set. Color represents the BH-adjusted p-value, and the gene ratio represents the number of overlapping DEG compared to the number of genes for a given term. (**D**) Correlation analysis between VS signatures and Kandoth et al. (2013) pan-cancer significantly mutated genes (SMG) mutational signatures.

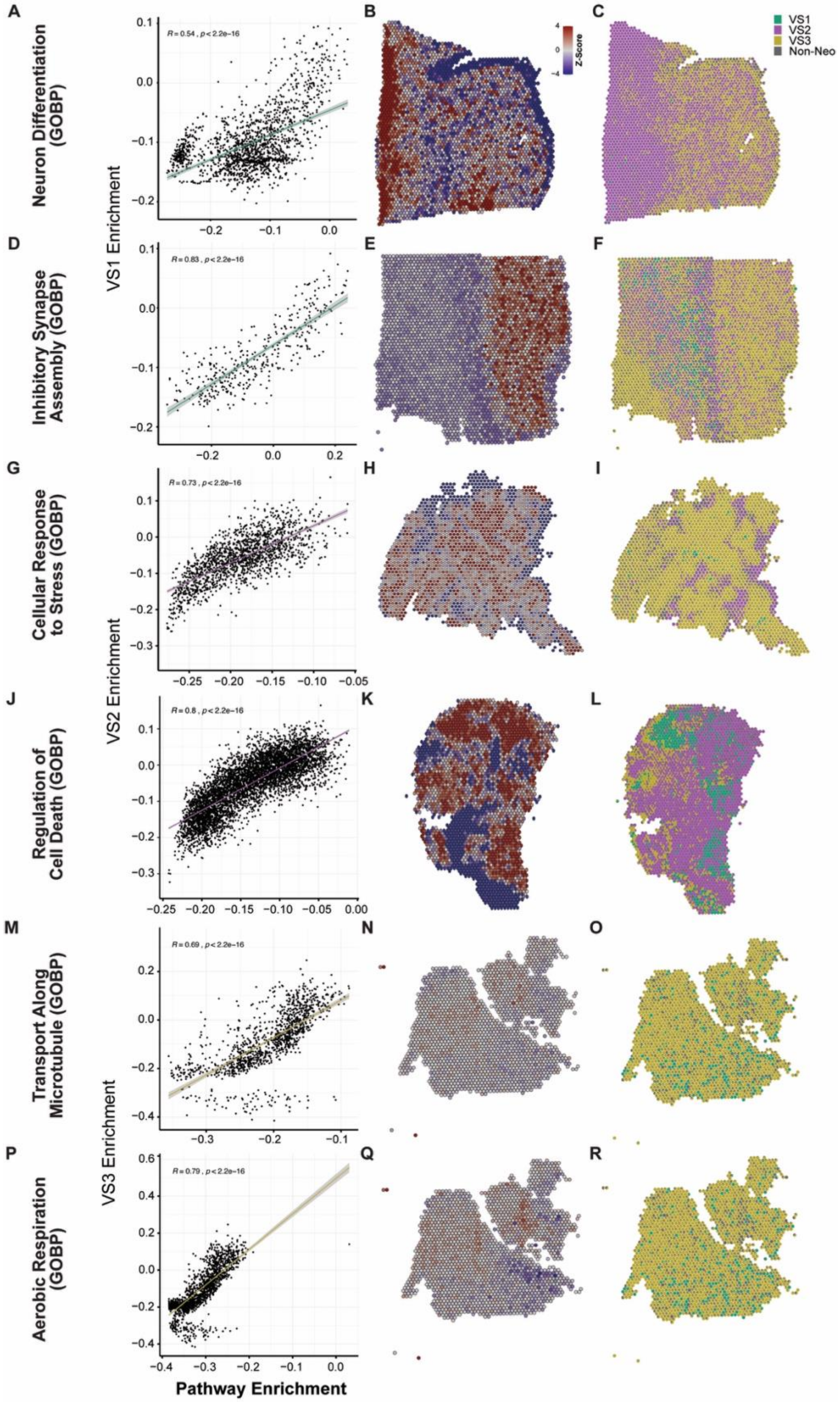

A
B
C
Neuron Differentiation (GOBP)
R = 0.54 , p < 2.2e-16
Z-Score
VS1
VS2
VS3
Non-Neo
D
E
F
Inhibitory Synapse Assembly (GOBP)
R = 0.83 , p < 2.2e-16
VS1 Enrichment
G
H
I
Cellular Response to Stress (GOBP)
R = 0.73 , p < 2.2e-16
J
K
L
Regulation of Cell Death (GOBP)
R = 0.8 , p < 2.2e-16
VS2 Enrichment
M
N
O
Transport Along Microtubule (GOBP)
R = 0.69 , p < 2.2e-16
P
Q
R
Aerobic Respiration (GOBP)
R = 0.79 , p < 2.2e-16
VS3 Enrichment
Pathway Enrichment

**Figure S8**. **Additional spatial transcriptomic spot-level correlation and enrichment analysis findings**

(**A, D, G, J, M, P**) Spot-level enrichment score scatter plot with Pearson's correlation analysis comparing VS singscore enrichment to the named GOBP pathway singscore enrichment. Each black dot represents a single spot with VS proportion > 50% by SCDC deconvolution (> 0% for VS1). Spots were also filtered to contain < 20% non-neoplastic cells to ensure the majority of cells in spots used for correlation analysis contain tumor cells. Colored line represents the line of best fit with error shaded in gray. (**B, E, H, K, N, Q**) Spatial scatter plot showing scaled GESECA enrichment scores for the named term (by row in the figure) at each spot. (**C, F, I, L, O, R**) Spatial scatter pie plots showing the proportion of each VS or non-neoplastic cells predicted to be present at each spot by SCDC deconvolution analysis.

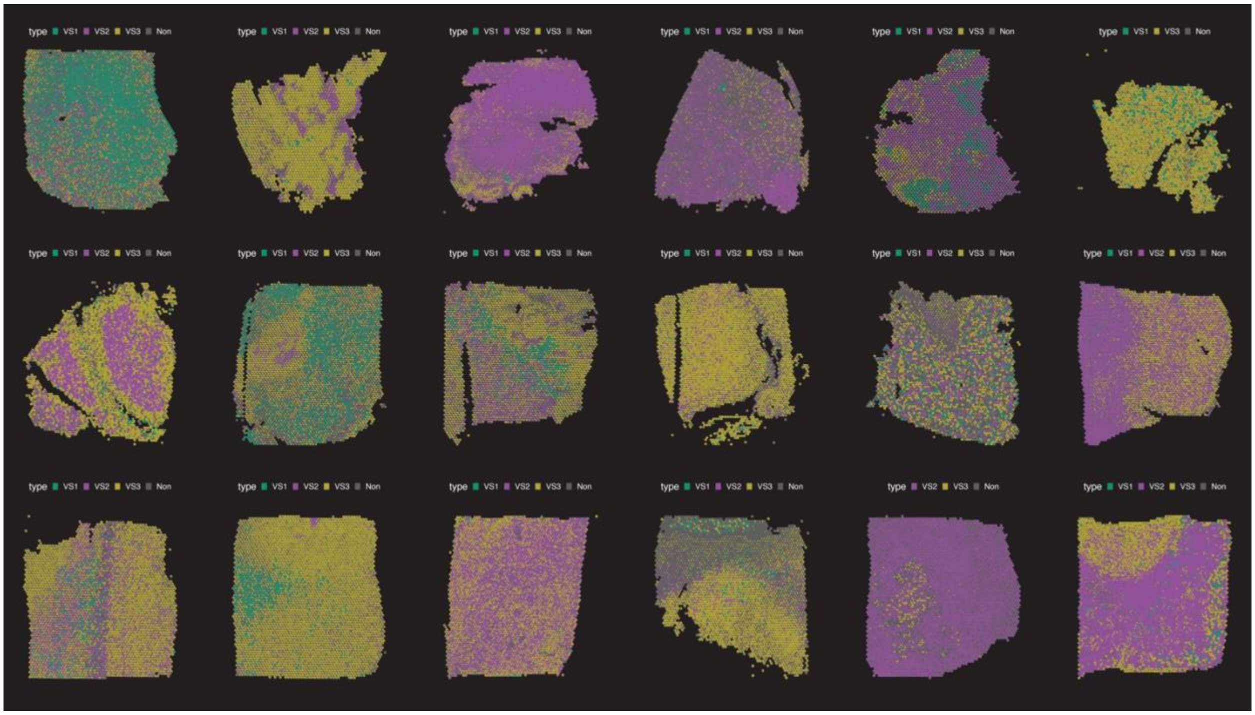


**Figure S9. Estimated vulnerability state proportions across 18 patient GBM sections profiled by stRNA-seq**

(**A-R**) Spatial scatter pie plots showing the estimated fraction of each VS or non-neoplastic cells predicted to be present at each spot by SCDC deconvolution analysis. Each 55 µm spot is a pie chart that represents the proportion of each cell type detected.

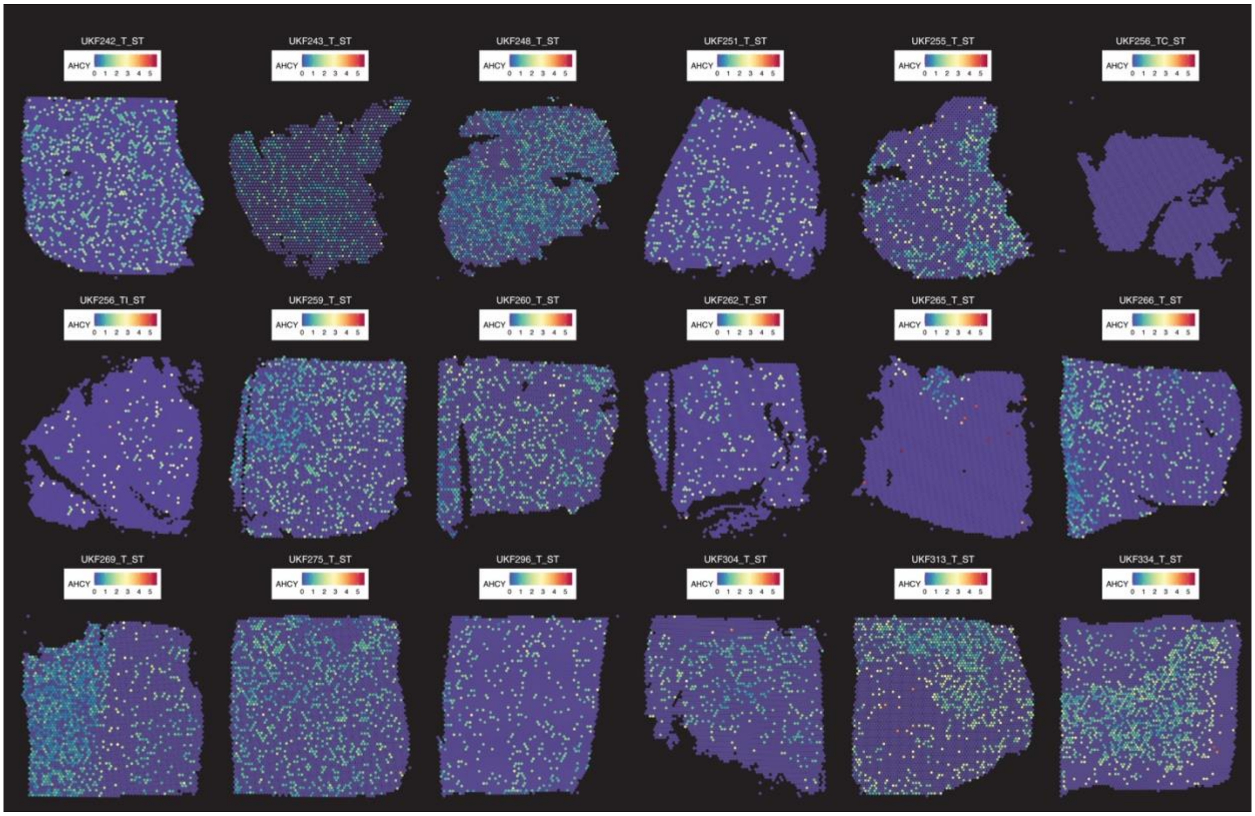


**Figure S10. Normalized AHCY expression in 18 patient GBM sections profiled by stRNA-seq**

(**A-R**) Spatial transcriptomic sections overlaid with normalized spot-level expression of AHCY detected by stRNA-seq at each 55 µm spot.

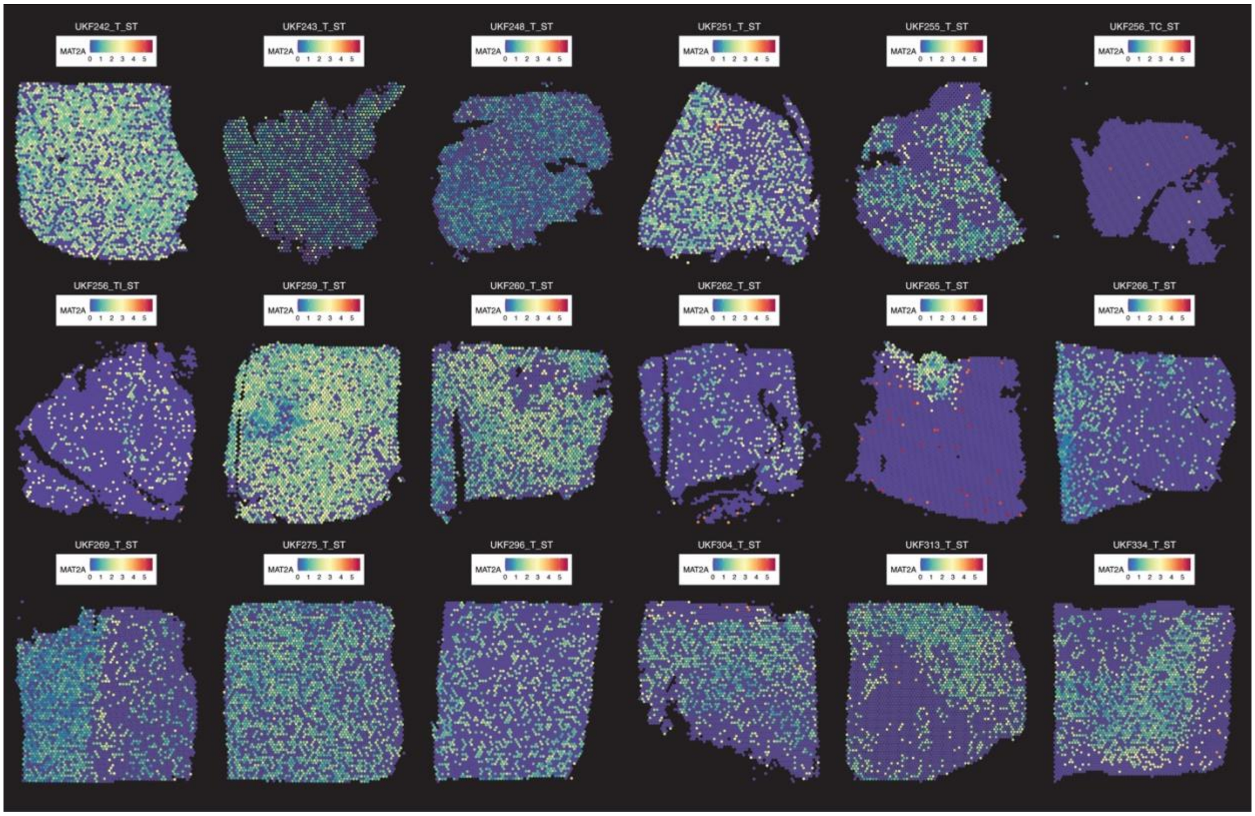


**Figure S11. Normalized MAT2A expression in 18 patient GBM sections profiled by stRNA-seq**

(**A-R**) Spatial transcriptomic sections overlaid with normalized spot-level expression of MAT2A detected by stRNA-seq at each spot.

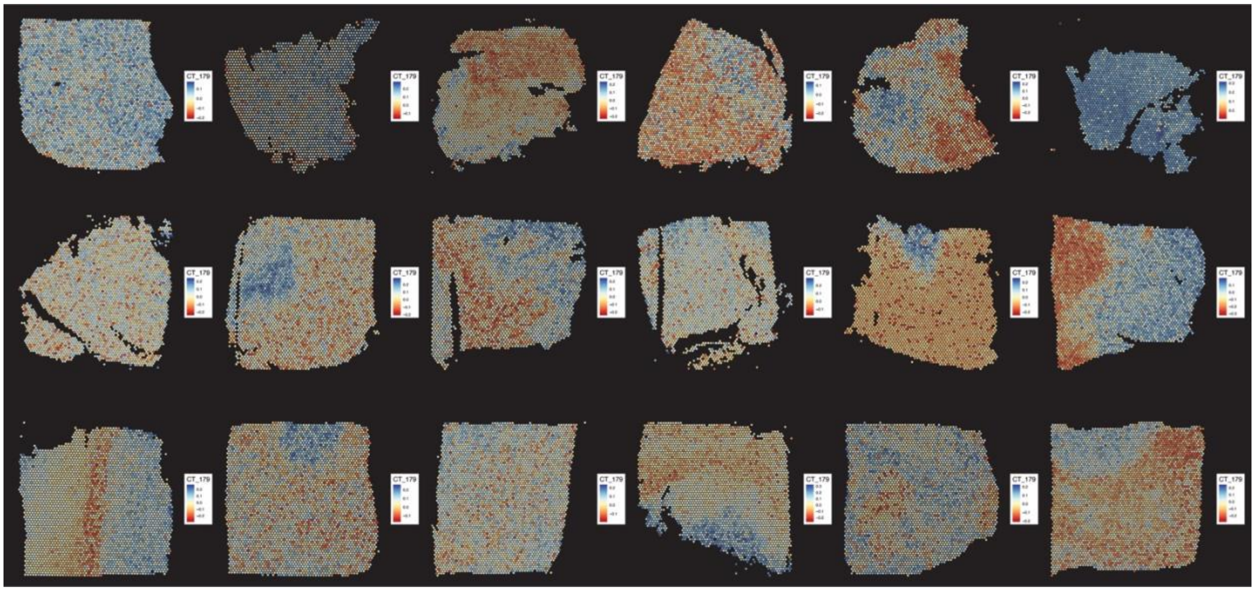

**Figure S12. CT-179 drug connectivity in 18 patient GBM sections profiled by stRNA-seq**

(**A-R**) Spatial transcriptomic sections overlaid with Fisher's Z-transformed CT-179 (OLIG2 inhibitor) drug connectivity scores. More negative scores (red) indicate greater spot-level drug sensitivity.

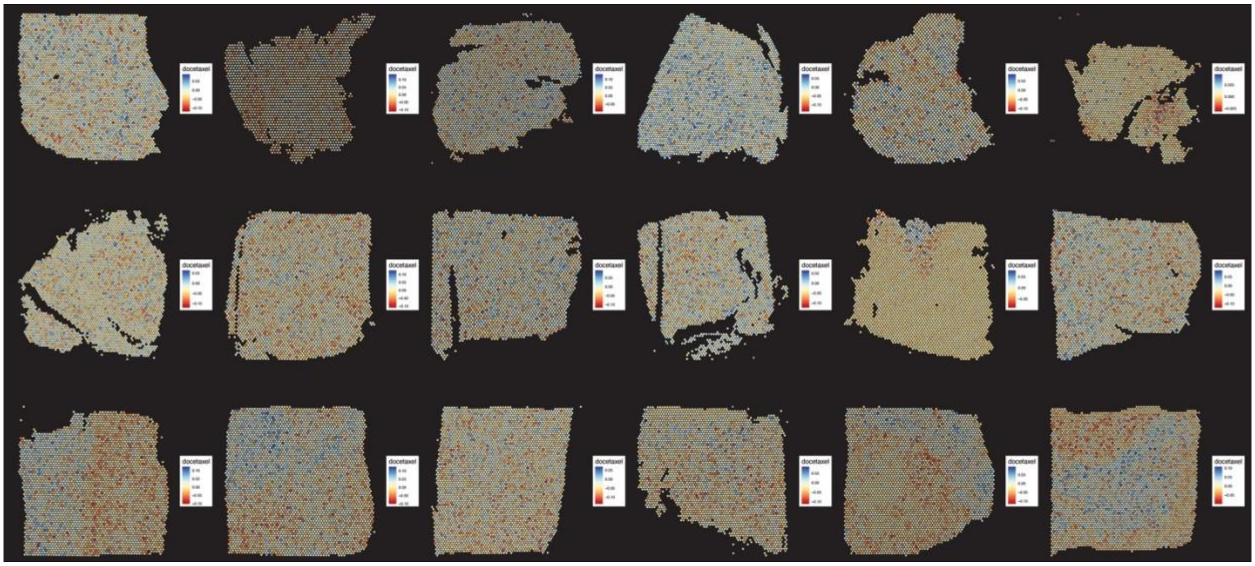

**Figure S13. Docetaxel drug connectivity in 18 patient GBM sections profiled by stRNA-seq**

(**A-R**) Spatial transcriptomic sections overlaid with Fisher's Z-transformed docetaxel (microtubule inhibitor) drug connectivity scores. More negative scores (red) indicate greater spot-level drug sensitivity.

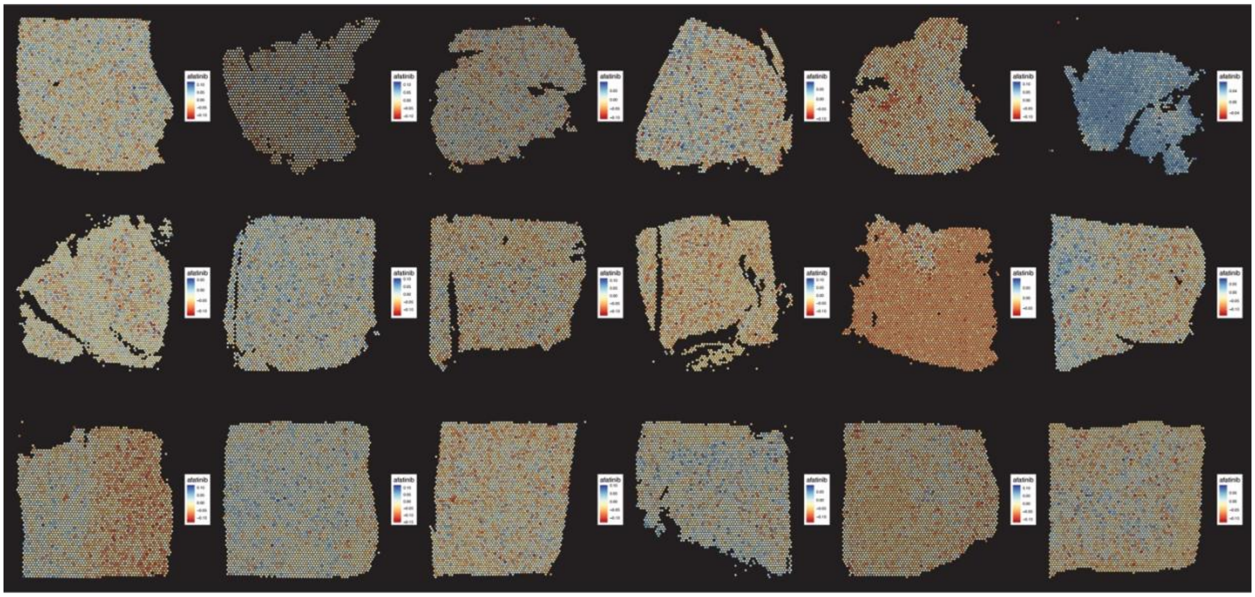

**Figure S14. Afatinib drug connectivity in 18 patient GBM sections profiled by stRNA-seq**

(**A-R**) Spatial transcriptomic sections overlaid with Fisher's Z-transformed afatinib (EGFR inhibitor) drug connectivity scores. More negative scores (red) indicate greater spot-level drug sensitivity.

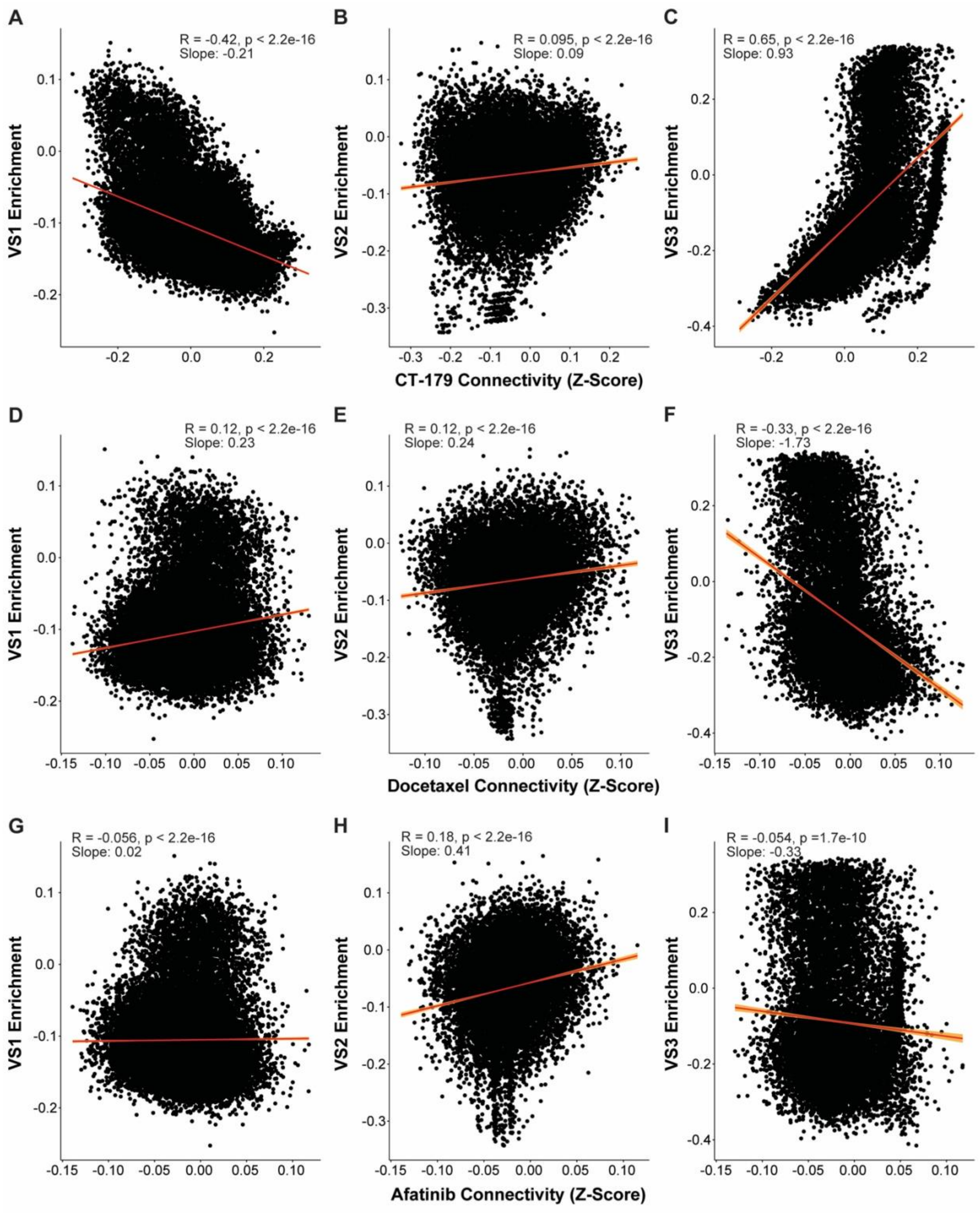

A
R = -0.42, p < 2.2e-16
Slope: -0.21
VS1 Enrichment
B
R = 0.095, p < 2.2e-16
Slope: 0.09
VS2 Enrichment
C
R = 0.65, p < 2.2e-16
Slope: 0.93
VS3 Enrichment
CT-179 Connectivity (Z-Score)
D
R = 0.12, p < 2.2e-16
Slope: 0.23
VS1 Enrichment
E
R = 0.12, p < 2.2e-16
Slope: 0.24
VS2 Enrichment
F
R = -0.33, p < 2.2e-16
Slope: -1.73
VS3 Enrichment
Docetaxel Connectivity (Z-Score)
G
R = -0.056, p < 2.2e-16
Slope: 0.02
VS1 Enrichment
H
R = 0.18, p < 2.2e-16
Slope: 0.41
VS2 Enrichment
I
R = -0.054, p =1.7e-10
Slope: -0.33
VS3 Enrichment
Afatinib Connectivity (Z-Score)

**Figure S15. Spatial VS enrichment correlates with spatial drug connectivity for OLIG2, microtubule, and EGFR inhibitors**

(**A-C**) Scatter plots showing spot-level VS enrichment vs. CT-179 Fisher's Z-transformed drug connectivity across all 18 10X Visium spatial tumor sections. (**D-F**) Scatter plots showing spot-level VS enrichment vs. docetaxel Fisher's Z-transformed drug connectivity. (**G-I**) Scatter plots showing spot-level VS enrichment vs. afatinib Fisher's Z-transformed drug connectivity. For all plots, red lines represent the line of best fit with correlation and statistical analysis performed using Pearson's R. The 95% confidence interval is indicated by the shaded orange region surrounding the line of best fit. The more negative the slope, the greater the predicted VS-specific sensitivity to a given compound.
